# Electrophysiological signatures of a developmental delay in a stem cell model of *KCNQ2* developmental and epileptic encephalopathy

**DOI:** 10.1101/2024.03.13.584717

**Authors:** Filip Rosa, Stephan Theiss, Susanne Krepp, Heidi Loeffler, Dulini Mendes, Stefanie Klingenstein, Stefan Liebau, Sarah Weckhuysen, Michael Alber, Steven Petrou, Holger Lerche, Snezana Maljevic, Thomas V Wuttke

**Author notes:** These authors contributed equally (Snezana Maljevic, Thomas V Wuttke). Corresponding authors: Snezana Maljevic https://orcid.org/0000-0003-1876-5872; Thomas V Wuttke https://orcid.org/0000-0001-5655-8490.

## Abstract

**Background:** *KCNQ2*, encoding K_V_7.2 ion channels, has emerged as one of the prominent genes causing early onset seizures with developmental delay (*KCNQ2* developmental and epileptic encephalopathy; *KCNQ2*-DEE). *KCNQ2 de novo* loss-of-function (LOF) and associated neuronal hyperexcitability have been accepted as mechanisms contributing to seizures. To investigate the developmental impact of *KCNQ2* LOF, we generated patient iPSC-derived models for two previously reported *de novo* variants, p.(Arg325Gly) and p.(Gly315Arg), linked to severe congenital DEE.

**Methods:** Functional investigation of the two variants was initially performed in *Xenopus laevis* oocyte system. Patient-derived iPSC lines were differentiated using NGN2- and embryoid body-based protocols yielding neurons roughly corresponding to mid- and mid-late gestational stages, respectively. K_V_7- mediated M-current, passive neuronal properties, action potential generation and spontaneous oscillatory network activities were analysed with whole-cell patch clamping.

**Findings:** Studied KCNQ2 variants showed LOF with a dominant-negative effect in the heterologous system. Reduced M-currents in patient iPSC-derived neurons corroborated a LOF as the main pathomechanism. Interestingly, this led to the reduced neuronal firing of the early neurons and to a disruption of complex oscillatory activity, with significantly reduced duration and amplitude of these events in patient iPSC-derived neurons.

**Interpretation:** We provide experimental evidence for changing roles of the M-current throughout development and place disease variant-mediated M-current reduction in the context of the neuronal maturation in the prenatal brain. Based on the reduced neuronal firing and disrupted oscillatory activity seen in patient iPSC-derived neurons, we propose that a delayed/impaired maturation of neuronal and network properties underlies *KCNQ*-DEE caused by LOF variants.

## Introduction

Developmental and epileptic encephalopathies (DEE) are a heterogenous group of neurogenetic disorders characterised by epileptic seizures, developmental delay (DD) and intellectual disability (ID). A subset of DEE with a specific clinical presentation, EEG and manifestation age are described as clinical syndromes (e.g. Ohtahara, West, and Lennox-Gastaut syndrome). With the discovery of many *de novo* or mosaic genetic variants as the cause of DEE, classification is increasingly based on the affected gene. A genetic aetiology can be confirmed in up to 50% of DEE cases with the number of affected genes reaching >900 (Oliver et al. 2023; Sánchez Fernández et al. 2019). Despite clinical heterogeneity, the majority of DEEs share an unfavourable prognosis, and selected treatments mainly focus on seizures without any impact on the neurodevelopmental symptoms.

*KCNQ2*, encoding the voltage gated potassium channel K_V_7.2, was one of the first genes linked to epilepsy. In a small number of families, variants in *KCNQ2* were identified as a cause of benign familial neonatal seizures, a self-limited epilepsy starting in the first days of life and remitting with phenobarbital or no treatment within couple of weeks to months (Biervert et al. 1998; Charlier et al. 1998; Singh et al. 1998). However, in the past ten years, *de novo* variants in this gene emerged as a leading genetic cause of early onset DEE or Ohtahara syndrome, frequently presenting with a burst suppression pattern in initial EEG recordings (Olson et al. 2017; Weckhuysen et al. 2012, 2013) followed by continued EEG changes including hypsarrhythmia (Kato et al. 2013; Millichap et al. 2016). *KCNQ2*-DEE usually apparent already during the first days of life, presents with frequent epileptic seizures accompanied by asphyxia, and progressively leads to profound DD and ID. Treatment is mainly based on sodium channel blockers, which lead to a significant seizure reduction but have no substantial impact on DD or ID (Berg et al. 2021; Pisano et al. 2015). Of note, a very recent retrospective clinical review of eight patients with *KCNQ2*-DEE suggested positive effects of the K_V_7 channel activator Retigabine on the seizure burden and also on the developmental outcome (Knight et al. 2023). However, Retigabine was withdrawn from the market in 2017 due to photoinstability and side effects thereby preventing further evaluations.

Homotetrameric K_V_7.2 and heterotetrameric K_V_7.2/K_V_7.3 channels are the main molecular correlates of the M-current, a subthreshold, slowly activating and deactivating, non-inactivating potassium current that can be blocked with muscarine receptor agonists. Very recent concepts consider the involvement of the M-current in different stages and processes of neurogenesis including regulation of proliferation of neural progenitors and establishing increasingly negative membrane potentials critical for progressive neuronal differentiation and functional maturation (Dirkx et al, 2020). Prior studies, however, largely focused on mature neurons and identified the role of the M-current in preventing hyperexcitability and enabling fine-tuning of action potential firing by controlling spike frequency adaptation. In this context, a reduced M-current would allow neurons to reach the action potential threshold faster thereby increasing neuronal firing, resulting in hyperexcitability and epileptic seizures. Indeed, *KCNQ2*-DEE variants predominantly cause a loss of channel function and a dominant-negative effect, implying that the mutant subunits also impair the function of the WT subunits in tetrameric channel complexes, was reported for a number of variants (Orhan et al. 2014; Zhang et al. 2020). Nevertheless, few gain of function variants with a less clear link to hyperexcitability have also been described (Mary et al. 2021; Miceli et al. 2015). Notably, 80% of KCNQ2-DEE pathological variants cluster in four regions of the K_V_7.2 protein. These domains are critical for the channel function and include the S4 voltage-sensor, the pore, the proximal C-terminal domain that binds phosphatidylinositol 4,5-bisphosphate (PIP2) and calmodulin (CaM A), and the more distal domain binding calmodulin (CaM B) (Millichap et al. 2016).

Disease modelling using human induced pluripotent stem cells has become an appealing approach to investigate pathophysiological mechanisms in the context of the specific genetic background of a studied patient. Neurodevelopmental disorders such as *KCNQ2*-DEE starting before or early after birth are probably well suited for such modelling, since the biological age of differentiated neurons that can be achieved *in vitro* can be estimated to correspond to foetal developmental stages (Rosa et al. 2020; Stein et al. 2014). We here generated several fibroblast- and keratinocyte derived iPSC lines from 2 patients with distinct LOF variants and associated *KCNQ2*-DEE as well as matched controls. Two neural induction protocols were used to generate 2D neuronal cultures: (i) forced expression of NGN2, a developmental transcriptional factor enforcing emergence of a rather uniform population of neurons (Busskamp et al. 2014; Ho et al. 2016; Zhang et al. 2013), and (ii) a small molecule embryoid body (EB) approach that will impact the pathways implicated in natural neurogenesis (Chambers et al. 2009; Zhang et al. 2001). These strategies yielded cultures with differential levels of developmental maturity, roughly corresponding to mid- to mid-late gestational stages, as reflected by intrinsic neuronal and action potential firing properties and the inability (NGN2 protocol) or ability (EB) to form complex spontaneous oscillatory network activities (Rosa et al., 2020).

In this study, we use these approaches to assess the impact of LOF *KCNQ2*-DEE variants on the early development and neuronal and network maturation.

## Results

Our study focuses on two previously reported *de novo KCNQ2* variants, p.(Arg325Gly) and p.(Gly315Arg) (in the following, we will use the one-letter code: R325G and G315R) (Weckhuysen et al. 2013), identified in two unrelated patients with *KCNQ2*-DEE. In both cases, epileptic seizures were already present on the 1^st^ day of life. Clinical presentation included asymmetrical/generalized tonic seizures accompanied by apnoea and desaturation. The initial EEG showed a burst-suppression pattern, with the follow-up revealing slow background activity. In the course of disease profound ID became apparent in both patients.

### R325G and G315R cause a loss of *KCNQ2*/Kv7.2 function

We first expressed R325G and G315R variants in *Xenopus laevis* oocytes and assessed the potassium currents using automated two-microelectrode voltage clamping (Orhan et al. 2014). When expressed as homotetramers, both variants caused a complete LOF, with currents staying at the background level (data not shown). The presumably predominant stoichiometry in heterozygotic patients was mimicked by co-expressing K_V_7.2-wildtype, K_V_7.2-mutant and K_V_7.3-wildtype subunits in a 1:1:2 ratio. In these experiments, the variants caused at least 50% reduction of the current amplitude (Kv7.2 + Kv7.3: 1.00 ± 0.05; R325G + Kv7.2 +Kv7.3: 0.48 ± 0.05; G315R + Kv7.2 + Kv7.3 0.31 ± 0.04; one-way ANOVA P<0.0001(*F*_2,58=56.72) with Dunnett‘s multiple comparison test: P(Kv7.2 + Kv7.3 vs G315R + Kv7.2 + Kv7.3)<0.0001; P(Kv7.2 + Kv7.3 vs R325G + Kv7.2 + Kv7.3)<0.0001; n(Kv7.2 + 7.3)=31, n(G315R+ Kv7.2 + Kv7.3)=17, n(R325G + Kv7.2 + Kv7.3)=13), which is larger than a 25% reduction expected for a single disrupted *KCNQ2* allele, confirming a LOF with a dominant-negative effect as the main mechanism for both variants (Fig 1B, C). Whereas G315R had not been analysed before, our data for R325G are in concordance with a previous characterization in CHO cells (Soldovieri et al. 2016).

**Figure 1.**
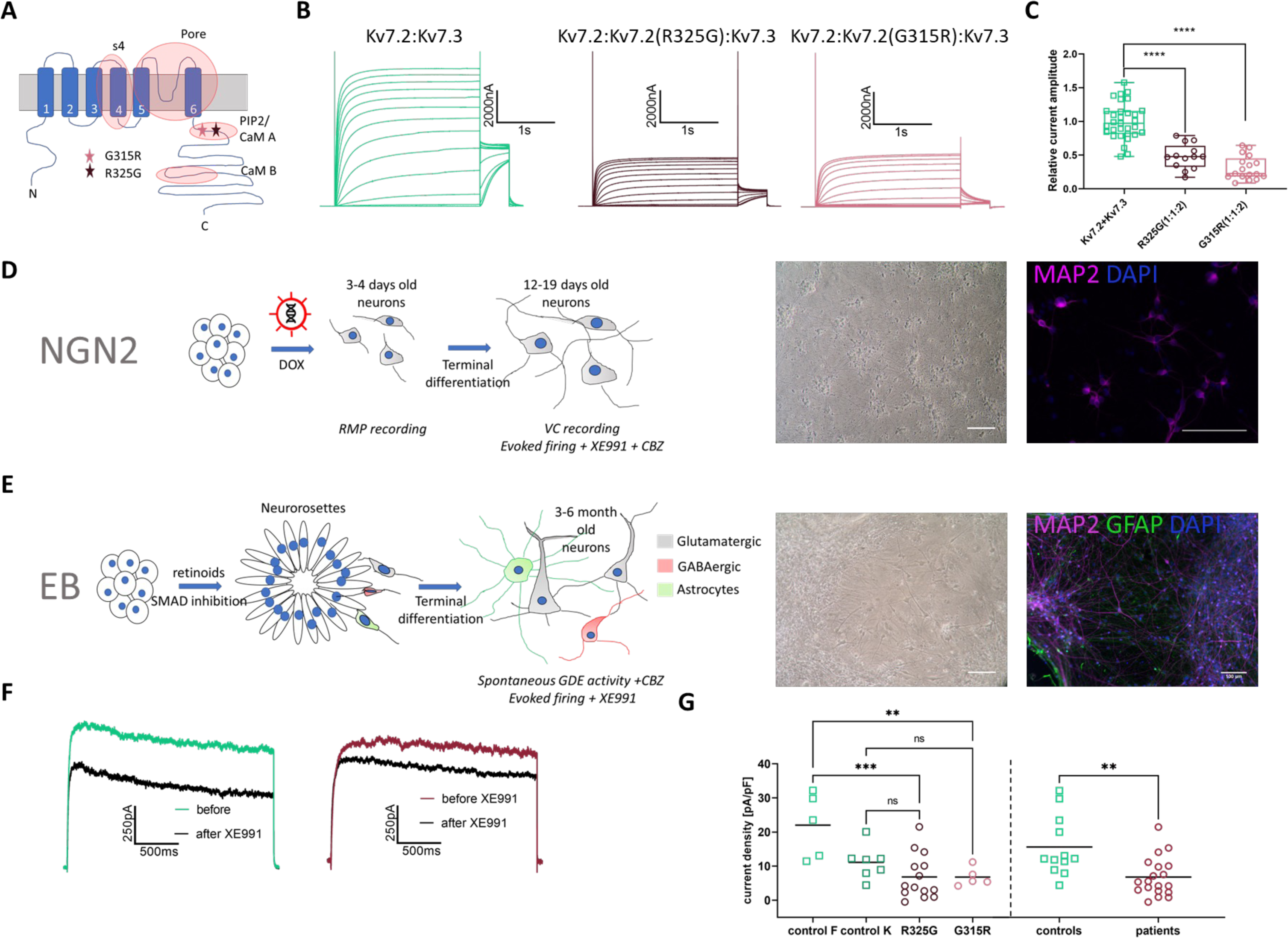
R325G and G315R are loss-of-function variants. (A) Kv7.2 tertiary structure with 4 hot spot zones harbouring 80 % of all KCNQ2 DEE variants. Positions of the two variants from this study are labelled with stars (modified according to Milichap, Park et al, Neurology 2016). (B) Representative current traces recorded from *Xenopus* oocytes expressing Kv7.2 + Kv7.3 heterotetramers. (C) Relative current amplitudes of heteromeric channels. All current amplitudes were normalised to the average current amplitude obtained from Kv7.2+Kv7.3 WT recorded on the same day; one-way ANOVA P<0.0001 with Dunnett‘s multiple comparison test: P(Kv7.2 + Kv7.3 vs G315R + Kv7.2 + Kv7.3)<0.0001; P(Kv7.2 + Kv7.3 vs R325G + Kv7.2 + Kv7.3)<0.0001; n(Kv7.2 + 7.3)=31, n(G315R+ Kv7.2 + Kv7.3)=17, n(R325G + Kv7.2 + Kv7.3)=13. (D) *left:* NGN2-differentiation workflow, *middle:* a bright field image of a 14-day old NGN2 culture, *right:* a representative image of MAP2 and DAPI staining of a 14-day old NGN2 culture. (E) *left:* EB-differentiation workflow, *middle:* a bright field image of a 6-month-old EB culture, *right:* a representative image showing MAP2, GFAP and DAPI staining of a 6-month-old EB culture. (F) Representative traces of M-current recorded at 25 mV in control (left) and patient-derived (right) neurons. (G) Scatter plot of the M-current density at the clamped voltage of 25 mV. Results for each control and patient line are shown on the left, pooled control and patient data on the right. For separate lines: One way ANOVA P = 0.0007, Holm-Sidak’s multiple comparison test: control F (fibroblasts) vs. R325G=0.0004; control F vs. G315R=0.0030; control K (keratinocytes) vs. R325G=0.4963; control K vs. G315R=0.6947; control F 22.01 ± 4.24 pA/pF, control K 11.10 ± 1.84 pA/pF, R325G 6.82 ± 1.74 pA/pF, G315R 6.80 ± 1.22 pA/pF; n(control F)=5, n(control K)=7, n(R325G)=14, n(G315R)=5; For merged genotypes: Unpaired T-test P=0.0019 controls: 15.65 ± 2.54 pa/pF, patients: 6.82 ± 1.31; n(control)=12, n(patient)=19; Scale bars 100µm; P<0.05: *; P< 0.01: **, P< 0.001: ***, P<0.0001: ****

### Patient iPSC-derived neurons are characterized by a reduced M-current

Following the analysis in *Xenopus laevis* oocytes, we generated induced pluripotent stem cell (iPSC) lines from both patients and two population controls. The iPSC lines were reprogrammed from patient fibroblasts (for R325G) or keratinocytes (for G315R), with corresponding population controls (Rosa et al. 2020). To compensate for the impact of the reprogramming procedure or potential somatic mutations, we generated 3 iPSC lines for the R325G (R325Ga,b,c) patient and 2 for the G315R (G315Ra,b) patient. Neural differentiation was performed using a fast protocol based on the overexpression of Neurogenin-2 (NGN2, Fig 1D) or an embryoid body protocol based on dual SMAD inhibition (Fig. 1E) (Rosa et al. 2020). In brief, the embryoid body protocol provides rather complex neuronal cultures containing glutamatergic and GABAergic neurons as well as glia cells in contrast to an almost uniform glutamatergic neuronal population and lack of glia cells by the NGN2 protocol. An exact cell allocation in the experiments can be found in Suppl. Fig. 2. A qualitative Western blot analysis confirmed K_V_7.2 expression in 12-19 days old NGN-2 neurons (human brain lysate served as positive control; Suppl. Fig. 1F). We performed patch clamp recordings of 12-19 days old neurons from both patients and controls and assessed the impact of the two variants on the M-current. The M-current was isolated by subsequent recordings in absence and presence of the M-current blocker XE991 (Fig. 1F depicts representative examples; see also Methods). Given that (i) the patient phenotypes are similar, (ii) the variants are only 10 amino acids apart and found in the same channel region, and (iii) have the same functional impact assessed in *Xenopus* oocytes, we also analysed pooled data for ‘variants vs. controls’. M-current densities are individually plotted for each patient and control (Fig. 1G, left panel), and are shown for each particular iPSC line in Suppl. Fig. 1G. Both variants caused a reduction of the M-current density of more than 50% (Fig. 1F, G) (for pooled genotypes: Unpaired T-test P=0.0019 controls: 15.65 ± 2.54 pA/pF patients: 6.82 ± 1.31 pA/pF; n(control pooled)=12, n(patients pooled)=19).

### Early depolarization block and depolarized resting membrane potential in patient-derived NGN2-neurons

The NGN2 protocol provides a uniform excitatory neuron population which can be traced roughly to the 2^nd^ trimester of human foetal development based on action potential firing and absence of synaptic activity (Rosa et al. 2020). Being characterized by a high input resistance of about 1,9 GOhm, the 14-19 days old NGN2 neurons function from the very beginning on the verge of a depolarization block, a silent state caused by reduced availability of voltage-gated sodium channels which are not released from their inactivated state. Using 800 ms-current steps, we observed that the patient neurons generated on average on the order of 4 action potentials (APs) upon 20 pA of injected current, whereas the control neurons produced approximately 7 APs upon the same stimulus (Fig. 2A). Furthermore, excitability was consistently reduced for each individual patient line in comparison to both control lines (Suppl. Fig. 3C, D). Interestingly, the M-current antagonist XE991 showed a comparable effect on the evoked firing in these neurons (Fig. 2B) supporting the notion that a functional M-current is required for the establishment of increasingly mature neuronal firing properties. There was no difference in the resting membrane potential between the 14–19-days old control- and patient-derived NGN2 neurons (Fig. 2C; individual lines Suppl. Fig. 3B). However, when comparing the resting membrane potential of less mature 3-4 days old neurons (characterized by very immature firing properties with generation of only a single action potential), we found that patient-derived neurons were depolarised by about 7mV in comparison to controls (For the pooled data: Mann-Whitney test P<0.0001 controls: -58.05± 0.84 mV, patients: -51.18 ± 0.92 mV; n(controls)=58 n(patients)=74) (Fig. 2D; individual lines Suppl. Fig. 3A). Altogether these data suggest that at early developmental stages functional M-current critically contributes to the establishment of gradually hyperpolarized resting membrane potentials of maturing neurons, required to release voltage-gated sodium channels from their inactivated state and eventually enabling action potential generation. At this point, a further rising M-current density provides a growing hyperpolarizing drive, ensuring efficient repolarization at the end of an action potential, finally enabling mature patterns of electrical excitability with generation of sustained trains of action potentials at various frequencies. *KCNQ2* LOF variants impair these early key functions of the M-current and result in a delayed progression of neurons from immature to mature functional properties, with likely ramifications also for downstream synaptogenesis and neuronal network formation.

**Figure 2.**
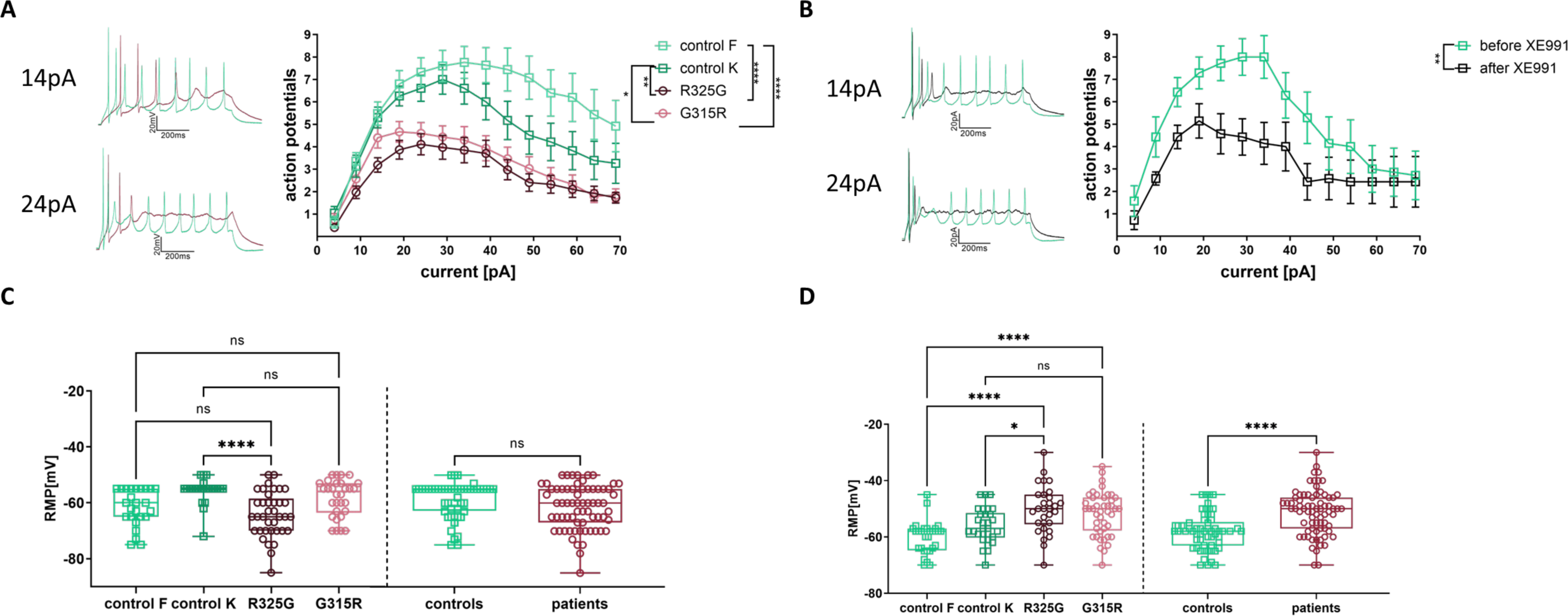
Characterisation of patient iPSC-derived NGN2 neurons. (A) Traces of evoked firing recordings obtained after 14 and 24 pA current injections for 800ms (left). Input-output curves showing the induced firing rates for control F, control K, R325G and G315R lines (right). Two-way repeated measurements ANOVA matched for cells. Genotype P<0.0001, n(control F)=25, n(control K)=23, n(R325G)=37, n(G315R)=30; Holm-Sidak’s multiple comparison test: control F vs. R325G<0.0001; control F vs. G315R<0.0001; control K vs. R325G=0.0030; control K vs. G315R=0.0336. (B) Traces of evoked firing recordings after 14 and 24 pA current injections for 800ms obtained from control NGN2 neurons before and after application of XE991 (left) and corresponding input-output curve (right). Two-way repeated measurements ANOVA matched for cells as well as for XE991 application. XE991 P=0.0040 n(cells)=7. (C) Resting membrane potential recorded in NGN2 differentiated neurons between days 14 and 19. Results for each separate patient and control line (left) and for the pooled control and patient lines (right). For separate lines: Kruskal-Wallis test P< 0.0001; Dunn’s multiple comparison test: control F vs. R325G> 0.9999; control F vs. G315R=0.2344; control K vs. R325G<0.0001; control K vs. G315R=0.7489; control F -61.64 ± 1.32 mV, control K -55.74 ± 0.95 mV, R325G -64.08 ± 1.31 mV, G315R -58.33 ± 1.20 mV; n(control F)=25, n(control K)=23, n(R325G)=37, n(G315R)=30. For the pooled data: Mann-Whitney test P=0.078 controls: -58.81± 0.92 mV, patients: -61.51 ± 0.96 mV; n(controls)= 48, n(patients)=67. (D) Resting membrane potential recorded in NGN2 differentiated neurons between days 3 and 4. Results for each separate patient and control line (left) and for the pooled control and patient lines (right). For separate lines: Kruskal-Wallis test P < 0.0001; Dunn’s multiple comparison test: control F vs. R325G< 0.0001; control F vs. G315R< 0.0001; control K vs. R325G<0.0261; control K vs. G315R=0.0791; control F -60.21 ± 1.12 mV, control K -56.03 ± 1.13 mV, R325G -50.30 ± 1.53mV, G315R -51.77 ± 1.15 mV. n(control F)=28, n(control K)=30, n(R325G)=30, n(G315R)=44. For the pooled data: Mann-Whitney test P<0.0001; controls: -58.05± 0.84 mV, patients: -51.18 ± 0.92 mV n(controls)= 58 n(patients)=74. P<0.05: *; P< 0.01: **, P< 0.001: ***, P<0.0001: ****

To test these hypotheses, we used a dual SMAD-inhibition embryoid body-based neural differentiation protocol to generate human patient- and control derived neurons with the ability to form synapses and spontaneous network activity reminiscent of physiological oscillatory neuronal activity found during foetal development.

### Magnitude and directionality of variant-mediated LOF effects depend on declining input resistance along maturation

We first performed neural differentiation of the patient (R325G) and control fibroblast iPS lines. In comparison to the NGN2 protocol, EB-based differentiation yielded neurons with lower (less immature) input resistances (on the order of 0,5 GOhm (EB derived neurons) versus 1,9 GOhm (NGN2 neurons)). Using 800 ms-lasting current injections to induce repetitive action potential firing of control neurons, we observed substantially more mature firing patterns (in comparison to the NGN2 protocol) with increased numbers of action potentials (Fig. 3A). In line with the decreased input resistance, injected current pulses needed to be overall higher than for NGN2-derived neurons (up to 120 pA versus 70 pA). Similar to the findings based on NGN2 differentiation, EB protocol-derived patient neurons were characterized by a reduced number of action potentials generated per current injection; although the magnitude of the difference was less pronounced than for the NGN2 protocol and varied between the individual lines (Fig 3A; individual lines Suppl. Fig. 3E). The effect of the genetic variant was again mimicked by recordings with the M-current blocker XE991 (Fig. 3B), pointing to the role of the M-current in providing hyperpolarizing drive for rapid termination of action potentials and for recovery of a sufficient proportion of voltage-gated sodium channels for the re-initiation of the subsequent AP. However, it is important to note that increasingly more mature neurons appear to rely less on K_V_7.2-mediated current for this particular task as indicated by milder XE991- and LOF-dependent (as mentioned above) effects on neurons derived by the EB versus the NGN2 protocol. This finding is in line with the shifting roles of the M-current, from an initial key factor for action potential generation in immature neurons towards a brake for repetitive firing in fully mature neurons (Dirkx et al. 2020).

**Figure 3.**
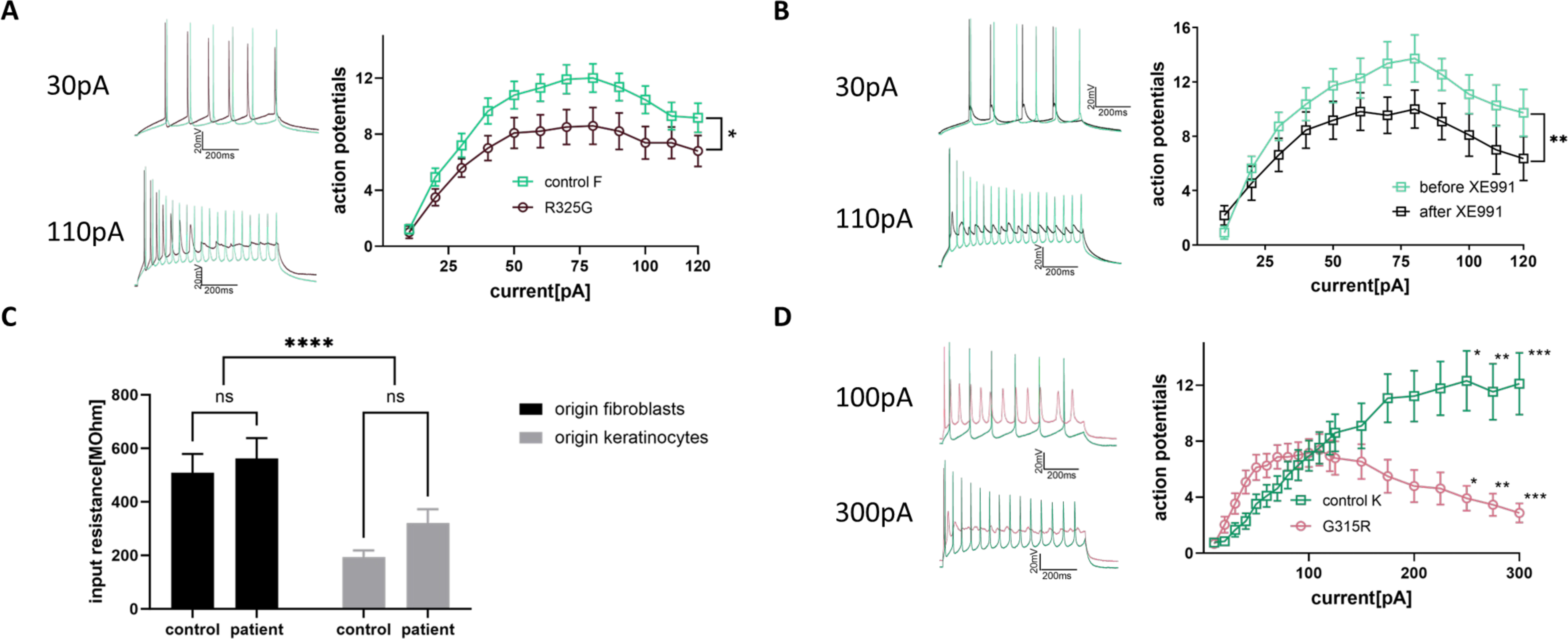
Characterisation of patient iPSC-derived EB neurons. (A) Traces of evoked firing recordings after 30 and 110 pA current injections for 800ms obtained from control and patient EB neurons with the fibroblast origin (left). Input-output curves for control F and R325G line (right). Two-way repeated measurements ANOVA matched for cells. Genotype P=0.017, n(control F)=31 n(R325G)=24. (B) Traces of evoked firing recordings after 30 and 110 pA current injections for 800ms obtained from control F EB neurons before and after XE991 application (left). Input-output curves for control before and after XE991 application (right). Two-way repeated measurements ANOVA matched for cells as well as for XE991 application. XE991 P=0.0032; n(cells)=11. (C) Comparison of input resistance between iPSC-derived neurons originating from different tissues. Two Way ANOVA tissue origin (fibroblast-versus keratinocyte derived) P<0.0001; mutation P=0.157; interaction P=0.561; neurons generated from fibroblasts: 532.6 ± 51.04 MOhm, neurons generated from keratinocytes 261.6 ± 30.83 MOhm; n(fibroblast-derived)=55, n(keratinocyte-derived)=45; Holm-Sidak’s test for control F vs. R325G P=0.9885; control K vs. G315R P=0.6897. (D) Traces of evoked firing recordings after 100 and 300 pA current injections for 800ms obtained from control and patient EB neurons with the keratinocyte origin (left). Input-output curves for control K and G315R line (right). Multiple unpaired t-tests with False Discovery Rate 1%, P(250 pA)=0.0012, P(275 pA)=0.0009, P(300pA)=0.0005; n(control K)=21, n(G315R)=24. All statistical testing was done with n(controls)=52 n(patients)=48 n(control F)=31, n(control K)=21, n(R325G)=24, n(G315R)=24. P<0.05: *; P< 0.01: **, P< 0.001: ***, P<0.0001: ****

Even though, fibroblast iPS line (EB protocol) derived neurons reached a higher level of maturity in comparison to NGN2-derived neurons (based on AP firing properties and input resistance), an input resistance of 533 ± 51 MOhm is still substantially different from mature primary mouse and bona fide human neurons with typical values around 150 MOhm (Schwarz et al. 2017, 2019; Testa-Silva et al. 2014). Given the postulated differential roles of the M-current during distinct developmental time windows (Dirkx et al. 2020), we next asked how *KCNQ2* variants would interfere with the excitability of human iPSC-derived neurons with substantially lower input resistances, comparable to values of primary mouse and human neurons.

In comparison to fibroblast derived neurons (EB protocol) we unexpectedly found that differentiation of the iPSC lines generated from keratinocytes indeed gave rise to neurons with profoundly lower input resistances (Two Way ANOVA original tissue P<0.0001; variant P=0.157; interaction P=0.561; fibroblast-derived neurons: 533 ± 51 Mohm, n = 55; keratinocyte-derived neurons 261.6 ± 30.8 MOhm, n = 45; Fig. 3C). In line with their reduced input resistance, current injections of up to 300 pA were tolerated and elicited trains of action potentials in control neurons without inducing depolarized plateaus and concomitant AP break-down (Fig. 3 D), which in turn increasingly occurred in neurons generated from fibroblasts starting from current injections of 80 pA onwards (Fig. 3A). Patient (G315R)-derived neurons revealed a bimodal activity pattern in comparison to control neurons, characterized by initial hyperexcitability (reflected by the generation of greater numbers of action potentials for current injections up to 110 pA), followed by a decline in firing rates at higher current injections (Fig. 3D; individual lines Suppl. Fig. 3F). Intriguingly, this firing pattern is highly reminiscent of recently published firing properties of postnatal P7 – P9 cortical layer II/III pyramidal neurons in a *KCNQ2*-DEE mouse model expressing the recurrent human dominant-negative variant p.T274M and particularly in control litter mates for recordings in presence of the M-current blocker XE991 (Biba-Maazou et al. 2022).

Taken together, these findings indicate that the function of the M-current strongly depends on the state of neuronal maturity and associated intrinsic properties. Consequently, LOF variants can convey quite opposite effects on neuronal excitability and therefore need to be interpreted in context of the maturational state of the analysed neurons. Furthermore, our data are in line with and support a previously introduced model of differential effects of the M-current during distinct stages of brain/neuron development (Dirkx et al. 2020).

### Giant depolarization events of patient-derived neurons are less developed

As we have previously shown (Rosa et al. 2020), iPSC-derived neurons using the embryoid body protocol develop early oscillatory activity, mainly driven by AMPA- and NMDA mediated glutamatergic events (Fig. 4 A-C). Whereas the low frequency (0-5Hz) activity is based on NMDA-R signalling (Fig. 4B), high frequency activity (10-100Hz) is related to AMPA-R signalling (Rosa et al. 2020) (Fig. 4C). This activity could present an *in vitro* analogy for spindle bursts or early network activity, i.e. the first synchronised activity in the developing brain which will be subsequently desynchronized by the increasing relative weight of GABAergic transmission (Garaschuk et al. 2000; Khazipov et al. 2004). In voltage-clamp recordings, giant depolarization events (GDE) at a clamped potential of -85mV are displayed as downward deviations of current (upper trace in Fig. 4 A), whereas in current-clamp recordings they occur as bursts of neuronal firing (lower trace in Fig. 4 A). The pooled patient-derived lines showed GDEs of shorter duration and smaller peak amplitude (Fig. 4D, E, F) than healthy controls (GDE duration: Mann-Whitney test P=0.0017; controls: 4.53 ± 0.23 s, patients 3.45 ± 0.23s, GDE peak amplitude: Mann-Whitney test P<0.0001; controls: 119 ± 14 pA, patient 61 ± 15 pA) and concordantly smaller areas under GDE (Fig. 4H, I). The area under the curve (specifically here the area under GDE) reflects current multiplied by time, thus the charge carried by a single GDE (calculated in pA.s (picoampere seconds) or pC (picocoulomb)) which is significantly smaller for GDEs generated by patient-derived neurons than control neurons (Mann-Whitney test P < 0.0001, controls 216 ± 32.8 pA.s, patients 73 ± 20.5 pA.s). Interestingly, the GDE events are more frequent in patient cells than in controls (Fig 4G) (Mann-Whitney test P=0.0406; controls: 1.933 ± 0.310 GDE/min, patients 2.858 ± 0.381 GDE/min); yet the total area under GDE normalised for one minute remains clearly larger in control lines (Fig. 4J) (Mann-Whitney test P < 0.0001, controls 1471 ± 327 pA.s/min, patients 335 ± 59 pA.s/min). Non-pooled control and patient line data are displayed in Suppl. Fig. 4 A-E.

**Figure 4.**
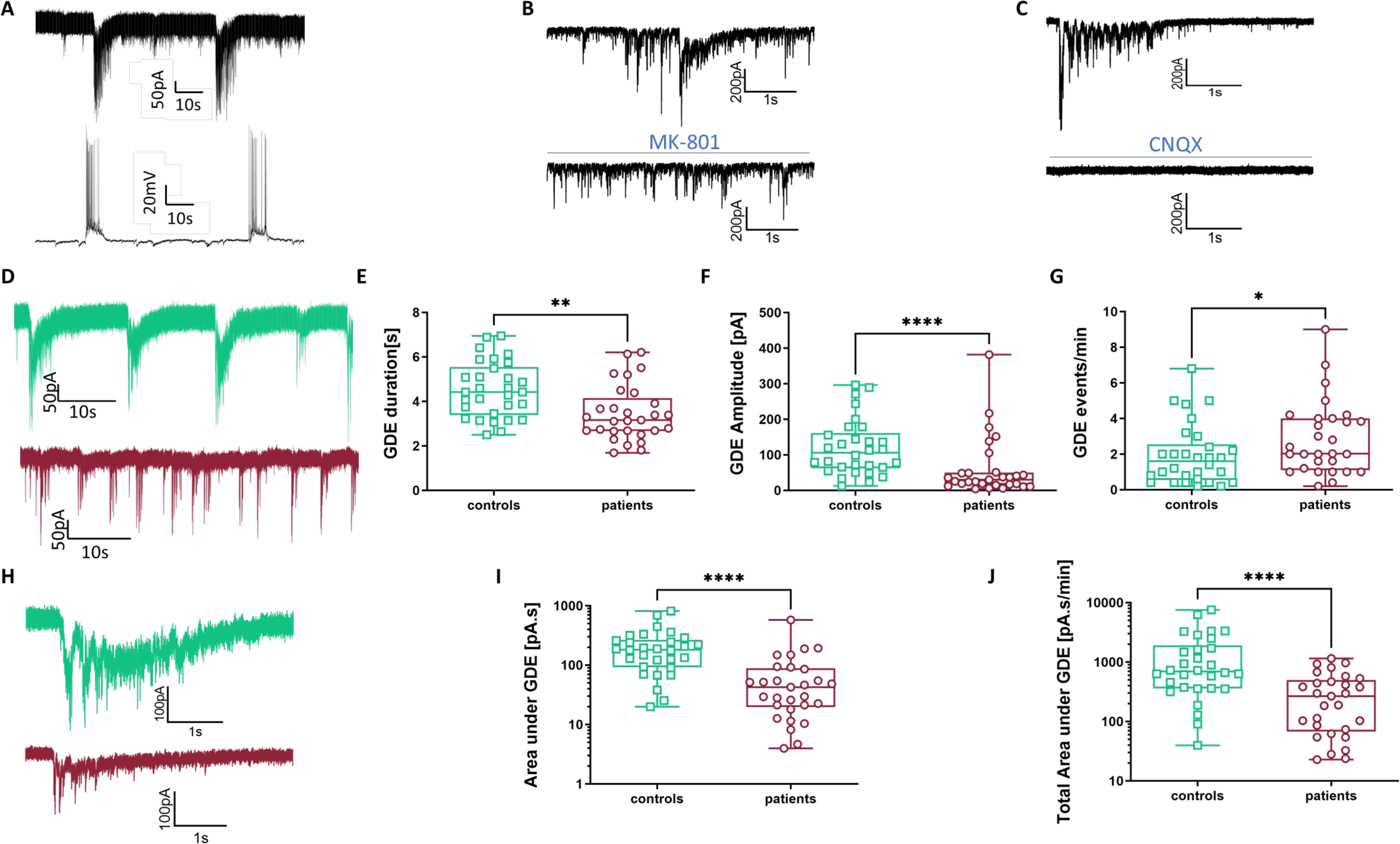
Patient iPSC-derived EB neurons generate smaller giant depolarization events (GDEs). (A) *top panel:* An example of a voltage clamp recording of GDE in a 5-month-old neuron (V=-85 mV); *bottom panel:* An example of a current clamp recording of GDE in the same 5-month-old neuron (RMP=-70mV). (B) Voltage clamp recording before (top) and after (bottom) application of NMDA-R blocker MK-801. (C) Voltage clamp recording before (top) and after (bottom) application of AMPA-R blocker CNQX. (D) Representative 1-minute traces of GDE recording at -85mV from control (top) and patient (bottom) iPSC-derived neurons. (E) GDE duration for pooled control and patient lines. Mann-Whitney test P=0.0017; controls: 4.53 ± 0.23 s, patients 3.45 ± 0.23s. (F) Peak amplitude of GDEs for pooled control and patient lines. Mann-Whitney test P<0.0001; controls: 119 ± 14 pA, patient 61 ± 15 pA (G) Number of GDE events in one minute for pooled control and patient lines. Mann-Whitney test P=0.0406; controls: 1.933 ± 0.310 GDE/min, patients 2.858 ± 0.381 GDE/min. (H) Representative traces of a single GDE event at -85mV from control (top) and patient (bottom) iPSC-derived neurons, (I) Area under an average GDE event for pooled control and patient lines. Y-axis logarithmic. Mann-Whitney test P < 0.0001, controls 216 ± 32.8 pA.s, patients 73 ± 20.5 pA.s (J) Total area under GDE event per one minute for pooled control and patient lines. Y-axis logarithmic. Mann-Whitney test P < 0.0001, controls 1471 ± 327 pA.s/min, patients 335 ± 59 pA.s/min. n(controls)=30, n(patients)=27. P<0.05: *; P< 0.01: **, P< 0.001: ***, P<0.0001: ****

Overall, these data demonstrate that patient-derived neurons exhibit less developed GDE activity compared to controls.

To further test a possible link between Kv7.2 function and the development of oscillations, we used primary hippocampal cultures derived from a knock-out (KO) *Kcnq2* murine model Neurons were obtained from E17 mice and neuronal activity was recorded using multielectrode arrays (MEA) on *day in vitro* (DIV) 10 and 24. We observed that the duration of bursting events recorded in cultures from WT mice increased by 4.8 fold within 14 days between DIV 10 and DIV 24, whereas cultures from homozygous KO mice showed only an increase by 1.8 fold in the same time span (WT 4.82 ± 1.33; KO 1.79 ± 0.37; n(WT)=12, n(KO)=10) (Suppl.Fig.4 F, G). These data obtained in an independent model system align well with the findings in human iPSC-derived neurons and collectively point toward the importance of Kv7.2 channels for the generation of oscillatory activity.

### Carbamazepine inhibits evoked firing and distorts GDE organisation

Carbamazepine (CBZ) is a well-known use-dependent blocker of sodium channels and has been reported to suppress seizures in *KCNQ2*-DEE and is used as first line treatment for early onset seizures (Millichap et al. 2016; Pisano et al. 2015; Sands et al. 2016). As previously suggested (Rosa et al. 2020), iPSC-derived immature neurons have not yet established appropriately hyperpolarized resting membrane potentials and lack sufficient hyperpolarizing drive to efficiently release voltage-gated sodium channels from the inactivated state and to enable firing of sustained trains of action potentials. Under these conditions a prominent effect of CBZ on neuronal firing can be expected. Indeed, 25 µM CBZ reduced the number of action potentials of 14-19 days old patient iPSC-derived NGN2 neurons by approximately 50% (Fig. 5A). The closer the cell was to a depolarization block, the more prominent was the effect. The trace recorded upon a current injection of 19pA (Fig. 5 left, upper panel) demonstrates the declining amplitude of action potentials due to an increasing fraction of inactivated sodium channels. Due to a reduced AP amplitude, not all potassium channels will get activated and consequently the hyperpolarization after an action potential will be reduced (Fig. 5A left, bottom panel), leading to a more positive membrane potential, which further supports sodium channel inactivation and facilitates CBZ binding to the channel. This may explain why a high efficacy of CBZ in immature neurons is further enhanced by decreased potassium conductances such as the M-current.

**Figure 5.**
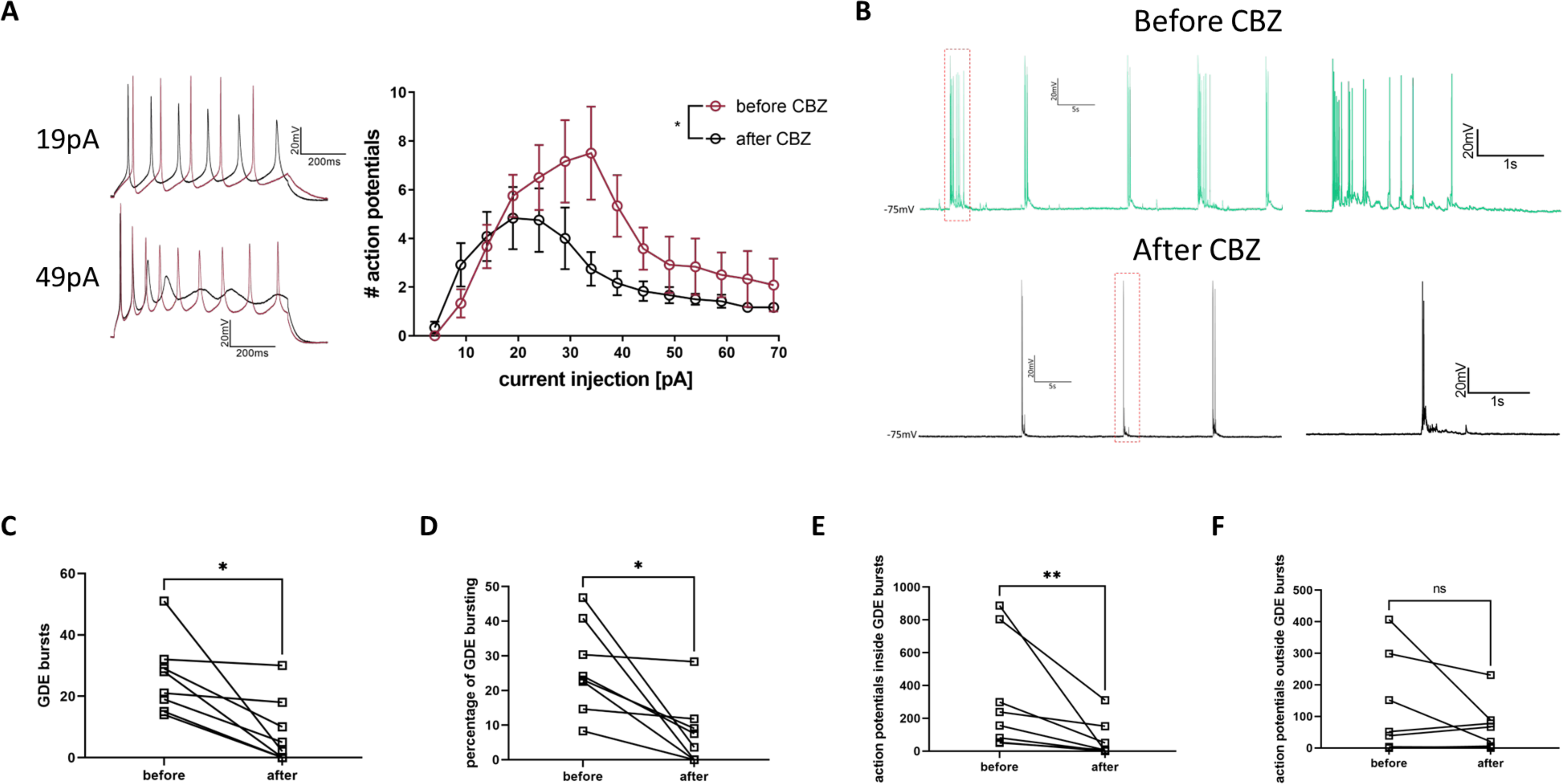
Carbamazepine (CBZ) deteriorates evoked firing in young neurons and perturbates GDE-bursting in more mature neurons. (A) Traces of evoked firing recordings for 800ms obtained from patient iPSC-derived NGN2 neurons (R325G and G315R) before and after 25µM CBZ treatment (left) and the corresponding input-output curves (right). Two-way repeated measurements ANOVA matched for cells as well as for CBZ application. CBZ P=0.0125; n(patients)=6. (B) Traces of spontaneous firing of control EB neurons before and after 25µM CBZ: 50s trace (left); magnified selected part of the trace (right). (C) Number of GDE bursts recorded during 5 min before and after CBZ application. Paired T-test P= 0.0117; before: 26.13 ± 4.253 bursts, after 8.13 ± 3.838 bursts. N(pairs)=8. (D) Percentage of GDE bursting recorded during 5 min before and after CBZ application. Paired T-test P= 0.0123; before: 26.45 ± 4.50 %, after 7.55± 3.88 %. (E) Number of action potentials inside GDE bursts recorded during 5 min before and after CBZ application. Wilcoxon matched-pairs signed rank test P= 0.0078; before: 321.3 ± 118.8 AP, after 65.75± 39.47 AP. (F) Number of action potentials outside GDE bursts recorded during 5 min before and after CBZ application. Wilcoxon matched-pairs signed rank test P= 0.4688 before: 119.1 ± 54.91 AP after 61.50± 27.34 AP.

To assess the effect of CBZ on the GDE activity, we performed recordings in control cells differentiated according to the EB protocol. CBZ (25 µM) suppressed the GDE bursting and relatively favoured single action potential firing (Fig. 5B), since single APs remained unaffected in the majority of cells (Supp. Fig. 6A shows a cell which completely changed its behaviour from GDE bursting to single action potential spiking). The GDE burst count declined within 5 minutes (Paired T-test P= 0.0117 before CBZ: 26.13 ± 4.25 bursts; after CBZ: 8.13 ± 3.84. N(pairs)=8) and the cumulative time of GDE bursting expressed as a percentage was reduced from 26% to 8 % (Paired T-test P= 0.0123; before: 26.45 ± 4.50 %, after 7.55± 3.88 %). The number of single action potentials outside GDE bursts was not affected by CBZ application over 5 min (Wilcoxon matched-pairs signed rank test P= 0.4688 before CBZ: 119.1 ± 54.9 AP, after CBZ: 61.5± 27.3 AP). However, the number of APs within bursts was profoundly reduced after CBZ application (Wilcoxon matched-pairs signed rank test P= 0.0078 before CBZ: 321.3 ± 118.8 AP; after CBZ 65.8± 39.5 AP), changing the ratio between action potentials inside and outside of bursts from 3:1 before to 1:1 after CBZ application. (Fig. 5C, D, E, F).

## Discussion

In this study, we investigated the impact of two dominant-negative LOF *KCNQ2* variants on the electrophysiological activity of human iPSC-derived neurons to elucidate early disease mechanisms of *KCNQ*-DEE. Neuronal cultures were differentiated from two patients with a severe clinical phenotype carrying two different de novo variants, p.R325G and p.G315R, respectively. iPSC-derived neurons were generated using two approaches (i.e. a fast NGN2-based differentiation protocol and a dual SMAD inhibition EB-based protocol) and yielded neurons of increasing levels of electrophysiological maturity. K_V_7.2 channel LOF translated into reduced M-current, depolarised resting membrane potentials and reduced action potential firing of NGN2 neurons. Strikingly, in 5-month-old neurons differentiated using a more complex EB-based protocol, both variants disrupted spontaneous complex oscillatory events (GDE). Figure 6 summarizes the results graphically in a developmental order. These findings are important as they suggest that *KCNQ2*-DEE variants interfere with the development of intrinsic and early oscillatory neuronal activity. Such activity patterns play important roles in circuit formation and regulation of developmental neuronal survival versus apoptosis, (Heck et al. 2008; Ikonomidou et al. 1999; Lebedeva et al. 2017) and their disruption may cause developmental delay and contribute to cognitive impairment as seen in patients with *KCNQ2*-DEE from the first days of life.

**Figure 6.**
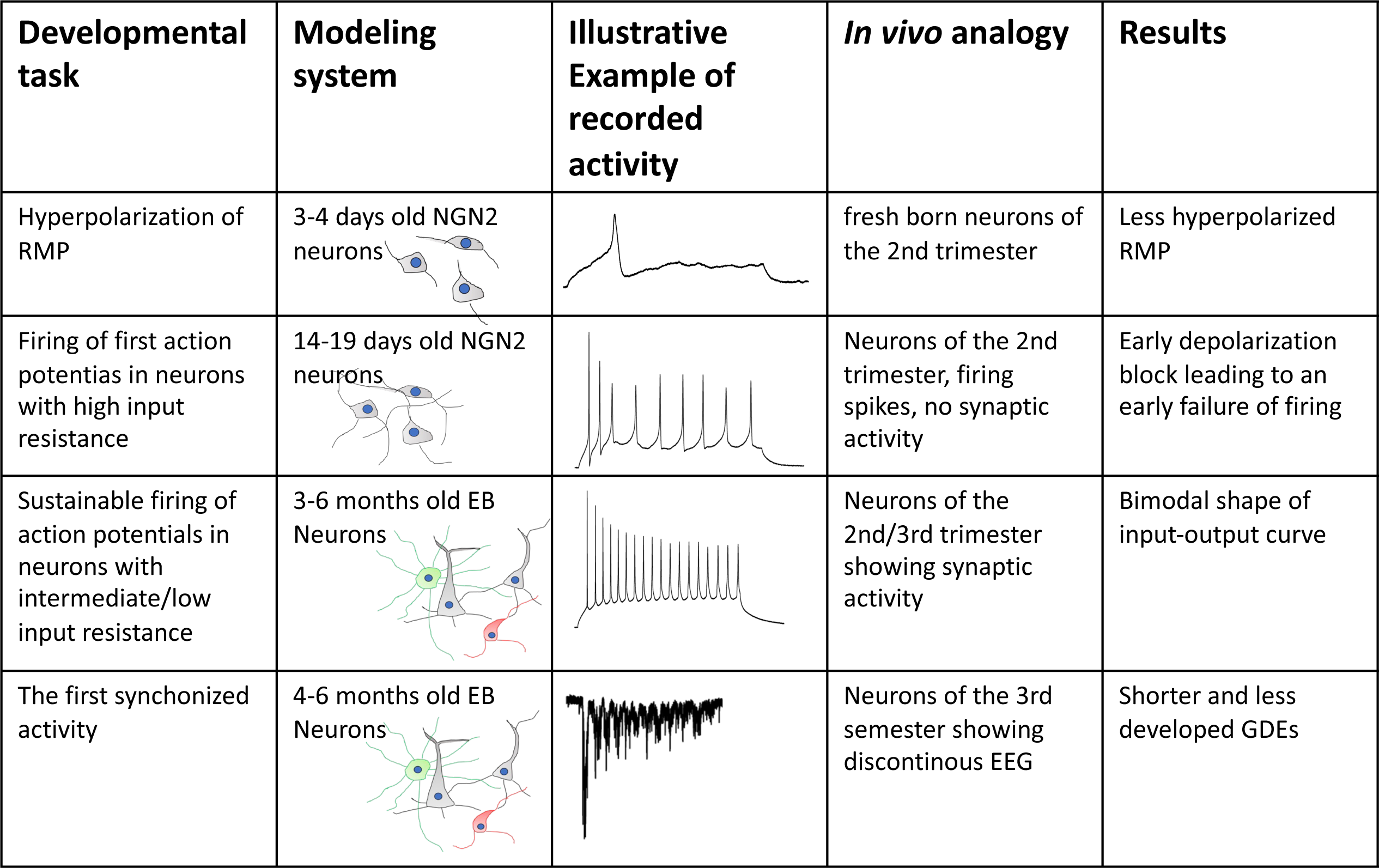
*KCNQ2*-DEE proposed developmental mechanisms. Developmental stages contributing to *KCNQ2*-DEE pathogenesis.

The very early expression of K_V_7.2 in human foetal development (Supp. Fig. 1E) is compatible with a role of the M-current in the gradual shift from depolarized membrane potentials of progenitor cells to hyperpolarized membrane potentials of mature neurons (Adams et al. 1982; Wladyka & Kunze 2006). The slowly activating non-inactivating low threshold potassium conductance is ideal for this developmental task. This was initially suggested by the work of Telezhkin et al. (2018) showing that a forced expression of Kv7.2 and Kv7.3 in iPSC-derived neurons accelerates their development. By reaching a membrane potential of -50mV, sodium channels can partially recover from inactivation and neurons start to fire action potentials. Our NGN2 iPSC data suggest that these processes are delayed in neurons derived from patients with *KCNQ2*-DEE due to impaired/ slowed progression from immature states characterized by depolarized resting membrane potentials, reduced hyperpolarizing drive, and precocious depolarization block. Blocking Kv7.2 by XE991 perfectly recapitulates this effect.

While the reduced firing seen in our study is in concordance with the reduced neuronal activity observed in the previously published KO iPSC model of *KCNQ2* (Deneault et al. 2018), it is in conflict with the data obtained in other studies and does not recapitulate the increased firing seen in murine models of Kv7.2 loss of function (Marguet et al. 2015; Niday et al. 2016; Peters et al. 2005; Soh et al. 2014). However, those experiments were commonly performed in more mature neurons. Indeed, we observed similar effects with our keratinocyte-based EB (dual SMAD inhibition) differentiation protocol, which yielded neurons of greater maturity and with concomitant lower input resistance in comparison to neurons generated from fibroblasts via dual SMAD inhibition or from fibroblasts/keratinocytes by NGN2 overexpression. Input-output (FI) curves of patient-keratinocyte-derived neurons (dual SMAD inhibition) showed a bimodal shape with generation of increased action potential numbers for low current stimulations which transitioned into decreased numbers for higher current injections. In fact this pattern is similar to the firing properties of postnatal P7 – P9 cortical layer II/III pyramidal neurons (which importantly were characterized by roughly comparable input resistances) in a *KCNQ2*-DEE mouse model expressing the dominant-negative *KCNQ2*-DEE variant p.T274M and particularly in control litter mates for recordings in presence of the M-current blocker XE991 (Biba-Maazou et al. 2022)).

It is essential to mention that iPSC-derived neurons model early developmental processes and thus the emergence and timing of neuronal activity patterns have to be interpreted within the framework of early developmental time windows and not based on terminology such as hypo-/ hyperexcitability or E/I (excitatory/inhibitory) balance as commonly done in models of epilepsy. In line with this notion Telezhkin et al. (2018) had demonstrated that forced expression of Kv7.2 and Kv7.3 in iPSC-derived neurons does not cause hyperexcitability but accelerates neuronal development/maturation as indicated among other parameters by a shortened time span to adopt increasingly higher action potential firing rates. Taking the level of electrophysiological maturity as a surrogate marker we deem *KCNQ2*-DEE patient iPSC-derived neurons less developed and not primarily hypoexcitable.

The last phase of prenatal development is characterised by low-frequency high amplitude oscillatory activity interrupted by long suppression phases which is called ‘*tracé discontinu’* (Khazipov & Luhmann 2006; Tolonen et al. 2007; Vanhatalo et al. 2005). Similar activity patterns, characterized by comparable event duration, frequency and involvement of neurotransmitters, were described in human neuronal stem cell-based cultures (Pruunsild et al. 2017; Trujillo et al. 2019; Zhong et al. 2017). In our previous study (Rosa et al. 2020), we named this activity “giant depolarization events” (GDE). GDE and GDE-like events present a first synchronized activity, an inevitable checkpoint, before a transition to postnatal brain activity and serve to enable proper wiring among neurons via calcium-flux, mediated by long depolarizations through positive feedback (Luhmann & Khazipov 2018; Luhmann et al. 2016; O’Leary et al. 2015; Takasu et al. 2002). Disrupted early oscillatory activity has the potential to interfere with the proper development of cortical architecture (Luhmann and Khazipov, 2018). A positive correlation between an abundance of slow oscillatory activity and brain growth in preterm babies (Benders et al. 2015) indeed suggests a possible clinical relevance of the presented human iPSC model-derived experimental data. IPSC-derived neurons from *KCNQ2*-DEE patients develop slow oscillatory activity (i.e. GDE), however less distinctive than neurons derived from healthy controls. These oscillatory events are mostly mediated by NMDA receptors (Rosa et al., 2020) which are highly permeable for Ca^2+^ and Na^+^ (Dingledine et al. 1999; Wollmuth & Sakmann 1998). GDEs are not only shorter in patient-derived neurons, but the overall charge (at least in parts carried by Ca^2+^ ions) moved across the neuronal membrane (total area under GDE) is four times smaller than in healthy controls. Ca^2+^ transients are critically involved in the regulation of gene expression, neuronal development, and maturation (Xin et al. 2005; Leinekugel et al. 1997; Wollmuth and Sakmann 1998; Minlebaev et al. 2009) and it is intriguing to speculate that subsequent maturation will be further hampered due to reduced Ca^2+^ influx. How a reduction of the M-current mechanistically leads to impaired GDE formation stays elusive. In principle the formation of GDE could either be simply delayed and could theoretically normalize over a prolonged time course (likely surpassing the feasible *in vitro* time span for viable neuron cultures) or a fully functional M-current could be vital for the proper formation of slow oscillations, thereby rendering the GDE impairment permanent.

Recent work points to a crucial role of the M-current in regulating cortical network dynamics in the context of wakefulness and slow wave sleep (Dalla Porta et al. 2023). Interestingly, the M-current is diurnally regulated by the cholinergic system and the phase of high M-current activity is associated with slow wave sleep (characterized by slow oscillations and high synchronicity). In contrast, the state of wakefulness (characterized by fast oscillations and desynchronisation) represents a phase of increased cholinergic inhibition and consequently of low M-current activity (Dalla Porta et al. 2023). These data may support the view that dysfunctional M-current probably leads to permanent deficits of various slow oscillatory activity patterns, likely not only in the context of foetal and early postnatal development but also with roles in regulation of transition from sleep to wakefulness, memory consolidation and cognition.

Carbamazepine presents a currently recommended anti-seizure medication for the treatment of infants with *KCNQ2*-DEE. It can reduce seizures by approximately 50% which is better than any other antiseizure medication (Kuersten et al. 2020; Pisano et al. 2015). However, no positive effect on developmental symptoms could be observed (Allen et al. 2014; Berg et al. 2021). In our study, the application of 25 µM CBZ on iPSC neurons derived from healthy controls had a negative impact on the burst characteristics during GDEs (recorded in current clamp). In general, GDE bursts were rarer, and the ratio between APs inside and outside bursts changed from 3:1 to 1:1. As discussed above, similar activities *in vivo* take important roles in neurodevelopment, indicating that CBZ could potentially interfere with such processes. Among all studied anti-seizure drugs taken by pregnant women with epilepsy CBZ had the most prominent impact on the brain size of the newborn (Almgren et al. 2009). It is tempting to speculate that the drug effects we observed on GDE in our *in vitro* studies could at least in parts translate to oscillatory activity patterns *in vivo*, such as SAT (spontaneous activity transients) (Tolonen et al. 2007), an activity pattern that has been shown to positively correlate with brain growth (Benders et al. 2015).

One limitation of our study is that it is based on only two patients. While we obtained comparable data for iPSC lines from both patients, the proposed concepts should be further validated in future studies and with additional patient samples. Another caveat is the absence of an isogenic control. However, multiple iPSC lines of the studied genotypes compensate partially for potential biases emerging from reprogramming and possible somatic mutations, whereas studying two different variants in the same protein segment compensates for genetic background. Finally, the specific K_V_7 blocker XE991 mimics the effects of the variants in healthy cells.

Recently, Dirkx et al. (2020) suggested that the expression of *KCNQ2* transcript variants may play a crucial role during early development and that repression of the M-current via a short dominant-negative transcript may enable proliferation and delay differentiation in early developmental stages. Indeed, we could show earlier (Rosa et al. 2020), that the short transcript is abundantly expressed in the pre-neuronal phases in cells derived from the EB protocol and harbours both studied variants. Since it is not expected that a short transcript conducts an ionic current (Hernandez et al. 2008; Schwake et al. 2003; Smith et al. 2001), we were not able to address this part of the development by electrophysiology. Therefore, the role of the short transcript in pathogenesis stays elusive.

In conclusion, our data indicate an early, prenatal onset of *KCNQ2*-DEE and suggest a slowed developmental pace with delayed neuronal maturation as initial pathophysiological mechanism. Future studies should increase the number of tested patient lines and assess the correlated gene expression alterations, which can corroborate the electrophysiological findings of this study.

## Methods

### Analysis in *Xenopus laevis* oocytes

*Site-directed mutagenesis to introduce the p.R325G and p.G315R variants* into *KCNQ2* cDNA in the pTLN vector

p.R325G and p.G315R variants were introduced into the wildtype *KCNQ2* cDNA cloned in the pTLN vector using the Quick Change PCR kit (Startagene) and verified by automated sequencing. Plasmid DNA was linearized using the Mlu1 restriction enzyme (Thermo Fisher) and then *in vitro* transcribed using the Sp6 RNA polymerase (Thermo Fisher), according to custom laboratory protocols.

#### Oocyte preparation and injection

*Xenopus laevis* oocyte extraction and use for recordings was approved by local authorities (Regierungspraesidium Tuebingen, Tuebingen, Germany). In the first step oocytes were treated by collagenase (1 mg/ml, CLS II collagenase, Biochrom KG, Berlin, Germany) in OR-2 solution (mM: 82,5 NaCl, 2,5 KCl, 1 MgCl2, 5 Hepes, pH 7,6), and subsequently washed and stored at 16°C in Barth solution (mM: 88 NaCl, 2,4 NaHCO3, 1 KCl, 0,33 Ca(NO3)2, 0,41 CaCl2, 0,82 MgSO4 ad Tris/HCL, pH7,4 with NaOH). To prevent the growth of bacteria 50ug/ml gentamicin (Biochrom KG, Germany) was added. cRNA concentrations were adjusted 0,8 μg/μl. The same amount of cRNA (50 – 70 nl) encoding WT or mutant K_V_7.2 or the combination with K_V_7.3 in a 1:1 ratio (K_V_7.2 WT / K_V_7.3 WT) or a 1:1:2 ratio (mimicking the presumed predominant stoichiometry in heterozygotic patients with one *KCNQ2* WT, one *KCNQ2* mutant and 2 *KCNQ3* WT alleles) was injected (Robooinject system; Multi Channel Systems, Reutlingen, Germany) in the same batch of oocytes plated in 96 well plates and measured at day 2-4 after injection.

#### Automated oocyte two-microelectrode voltage clamp

The currents were recorded at room temperature (20-22 °C) using roboocyte2 (Multi Channels Systems, Reutlingen, Germany). The intracellular glass microelectrodes had a resistance of 0.3 - 1 MOhm filled with 1M KCl/1.5 KAc. As bath solution we used ND96 (in mM: 93.5 NaCl, 2 KCl, 1.8 CaCl2, 2 MgCl2 and 5 Hepes; pH 7.5). The sampling rate was 1 kHz. Recordings were performed from a holding potential of -80 mV and 2 s-long depolarising steps from -80 to +50 mV in 10mV increments followed by a 0.5 s-long -30 mV step to record tail currents were applied. Amplitudes were analysed (Roboocyte2+ software; Multi Channel Systems, Reutlingen, Germany) at the end of the pulse to +40 mV and normalised to the mean of the WT recorded on the same day so that the data could be pooled together.

### iPS cell models

#### Generation of iPS cells

iPSC lines were generated from fibroblasts (R325G, 3 lines) or keratinocytes (G315R, 2 lines). Two corresponding control lines, Control F and Control K were generated from fibroblasts (F) or keratinocytes (K) obtained from healthy individuals (Rosa et al., 2020).

After establishing the primary cultures of fibroblasts and keratinocytes, the cells were reprogrammed by a mixture of retroviral vectors encoding OCT3/4 (pMXs-hOCT3/4 Addgene: 17217), SOX2 (pMXs-hSOX2 Addgene: 17218), KLF4 (pMXs-hKLF4 Addgene: 17219) and MYC (pMXs-hc-MYC Addgene: 17220) and subsequently frozen after a quality control which included (a) validation of the silencing of viral transgenes, (ii) aberration-free karyotype and (iii) examination of expression of the markers of pluripotency using immunostaining. A detailed description can be found in Rosa et al. (2020).

#### NGN2-based neural differentiation

120 000 iPS cells were plated on 12-well plates one day before the transduction by lentiviruses pLV_TRET_hNgn2_UBC_Puro (Addgene: 61474) and pLV_hEF1a-rtTA3 (Addgene:61472) as described in Rosa et al. (2020). For a selection of successfully transduced cells puromycin (Invivogen, USA) in the concentration of 1 µg/ml was added to E8 culturing medium (Thermo Fisher, USA). After 2 days cells were manually passaged and the successfully transduced lines were established, aliquoted and frozen.

The NGN2-driven neurodifferentiation was promoted by adding 2.5 µg/ml doxycycline (Sigma, USA) for 4 hours followed by dissociation by Accutase (Capricorn, DE) and plating 80000 cells per coverslip in NSC Medium. For the next 5 days the cells were kept in NSC Medium containing 1 µg/ml doxycycline. The detailed protocol can be found in Rosa et al. (2020). Voltage clamp experiments were performed between days 12-19, RMP was recorded between days 3-4 and 14-19. The firing properties of NGN2-differentiated neurons were recorded between days 14 and 19.

#### EB-based neural differentiation

Large pieces of similar size (diameter 200 µm) were dissected manually from iPSC colonies growing in mTeSR Medium (Stemcell Technologies, CA). The pieces were let form embryoid bodies over 5 days. To support a neurodifferentiation, the NSC medium supplemented with 10 ng/ml FGF2 (Peprotech, USA) was used. On day 5, the embryoid bodies were allowed to adhere to matrigel-coated (Corning, USA) plates to form neurorosettes. To enhance their neuronal fate NSC medium was enriched by 1 µM dorsomorphin (Stemgent, USA) and 10 µM SB-431542 (Tocris, UK). The selected neurorosettes were allowed to form neurospheres in T25 flasks. The cultivation continued for the next 7 days in NSC medium containing 10 ng/ml FGF2. On day 21, the terminal differentiation was initiated by triturating neurospheres to approximately 200 µm large pieces, which adhered to coverslips coated with poly-L-ornithine and laminin (both Sigma, USA). The ongoing differentiation continued in house made BrainPhys Medium (Rosa et al. 2020), supplemented with 10 ng/ml BDNF, 10 ng/ml GDNF, 10 ng/ml IGF1 (all 3 Peprotech, USA), 200 µM ascorbic acid, 1µM cAMP, 1 µg/ml laminin (all 3 Sigma, USA). During terminal differentiation, the media was changed weekly. After the first month, no supplements were added. Assessment of action potential firing was performed on 3-6 months old EB-derived neurons. A detailed description can be found in Rosa et al. (2020).

#### Electrophysiological recordings

Whole cell patch clamp recordings were performed at 22°C. For all experiments same extracellular (in mM: 140 NaCl, 4.2 KCl, 1.1 CaCl_2_, 1.0 MgSO_4_, 0.5 Na_2_HPO_4_, 0.45 NaH_2_PO_4_, 5 HEPES, 10 glucose pH: 7.4) and intracellular solutions (in mM: 135 K-gluconate, 4 NaCl, 0.5 CaCl_2_, 10 HEPES, 5 EGTA, 2 Mg-ATP, 0.4 Na-GTP, pH 7.3) were used. The electrophysiology rig was equipped with an Axopatch 200B (Molecular Devices, USA) amplifier connected to a Digidata 1440A (Molecular Devices, USA) and running pClamp10 (Molecular Devices, USA) acquisition software. Perfusion was maintained by a Gilson peristaltic pump.

After establishing a giga-seal, the cells were controlled for their quality. Only neurons with RMP<-50 and access resistance < 20MOhm were accepted for further experiments. For voltage clamp recordings, currents were sampled at 20 kHz and lowpass filtered below 5 kHz. For the M-current voltage clamp experiments, the protocol included 2.5 s long depolarising steps starting from -75mV to 25mV in 20mV increments. After the baseline recording, cells were perfused by the same extracellular solution supplemented by 10µM XE991(M-current blocker: Sigma) for 5 minutes. During this perfusion time the capacitance, series resistance and access resistance were checked in 2min intervals. Only cells with <20% change in the original values were accepted for the analysis. M-current density was calculated as a difference between a current recorded at +25mV before and after the addition of XE911 divided by the cell capacitance. Current amplitude was determined as the mean value of the current recorded between 2000 and 2500 ms. To block sodium currents and A-type potassium currents, the extracellular solution contained 1 µM TTX (Alomone, IS) and 4mM 4-AP (Sigma, USA), respectively.

For the current clamp experiments, the sampling rate was 100kHz with a low pass filter set below 10kHz. The evoked firing of NGN2-differentiated cells was recorded using a protocol with 15 steps (800ms). The initial current injection was -1pA and increments of 5pA were applied. Evoked firing of EB-differentiated neurons was recorded using a protocol with 13 steps (800ms). The initial current injection was 0pA and increments of 10pA were applied. For keratinocytes-derived lines a depolarization block could not be reached with this protocol; for these lines, a protocol starting at a current injection of 125pA with incremental steps of 25pA to reach a maximal injection of 300 pA was used. To determine input resistance, we injected four 500 ms-long current steps from -10 pA with an increment of -10 pA.

To block M-current we used 10µM XE991 and to demonstrate the effect of carbamazepine we used 25 µM CBZ (both Sigma, USA).

To record giant depolarization events, neurons were clamped at -85 mV (reversal potential for Cl^-^) for 5 min with a sampling rate of 100 kHz and a 10 kHz lowpass filter. Cells which could not be recorded for the entire 5 min were included in the analysis as long as they fulfilled the quality criteria (RMP and series and access resistance not deteriorated by more than 20%). The time-dependant parameters were normalized per one minute.

Recordings with CBZ were performed in current clamp mode: First, a 5-minute baseline recording of spontaneous activity was performed without CBZ. Afterwards, perfusion with an extracellular solution containing 25 µM CBZ was started. 5 minutes were allowed for equilibration and then followed by a 5-minute recording in presence of CBZ.

#### Immunocytochemistry

Cells were plated on coverslips coated with PDL (Sigma, USA). After removing the media, washing steps in PBS containing Ca^2+^ and Mg^2+^ preceded the fixation in 4 % formaldehyde for 15 min at room temperature. For blocking, cells were incubated for 1 h at room temperature in TBS (50 mM Tris-Cl, pH 7.6, 150 mM NaCl) with 10 % normal goat serum and 0.2 % Triton. All primary antibodies were incubated overnight in 1:20 diluted blocking solution at 4°C. Secondary antibodies were diluted 1:500 in TBS and left on cells for one hour at room temperature. Nuclei were stained by 4’,6-Diamidine-2’-phenylindole dihydrochloride (DAPI) (Sigma, USA) in 1:5000 dilution applied for 2 min at room temperature. 3-4 washes with TBS were performed between each step and coverslips were rinsed with water before mounting with Fluoromount-G™ Mounting Medium (ThermoFisher, USA) on a microscope slide.

Epifluorescence imaging was performed on a Zeiss Axioimager M1 microscope. Fiji software was used for exporting and editing the images.

The following primary and secondary antibodies were used: antiMAP2 (chicken, 1:2000, Abcam, UK); antiGFAP (rabbit, 1:500, Dako, USA), antiOCT4 (rabbit, 1:500, Abcam, UK), antiTRA1-60 (mouse, 1:500, Abcam, UK).

#### RT-qPCR

For RNA isolation, we used the ISOLATE II RNA mini kit including on-column DNA digestion (Bioline, UK). The concentrations were measured by NanoDrop ND-1000 (Peqlab, DE). To test silencing of viral transgenes and upregulation of stem cell markers, we used the SensiFAST SYBR Lo-ROX one step kit (Bioline, UK) and the ΔΔCt method with GAPDH serving as a housekeeping gene (Reinhardt et al. 2013).

#### Western blot

Cells were lysed using the following buffer (in mM): 20 Tris (pH 7.5), 150 NaCl, 1 EDTA, 1 EGTA, 2.5 Napyrophosphate, 1 β-glycerolphosphate, 1 sodium-orthovanadate, 10 DTT, 1% Triton and 1x cOmplete solution (Roche). Total protein concentration was measured via Bradford assay. 12% polyacrylamid gels were loaded with 20µg of total protein per lane to separate these by sodium dodecyl sulfate-polyacrylamide gel electrophoresis (SDS PAGE). After transferring the proteins onto a nitrocellulose membrane (Whatman) via electrophoresis at 4°C in Towbin buffer (25 mM Tris, 192 mM glycine, pH 8.3, 10% (v/v) methanol), the blots were blocked in 5% non-fat dry milk powder in phosphate-bufferd saline with 1% Tween (PBST) for 1h at RT. Subsequently, membranes were probed with a polyclonal rabbit primary antibody against K_V_7.2 (Pineda) at 1:1000 overnight at 4°C. Following this, the membranes were washed and shaken in PBST thrice and then re-probed with a secondary goat anti-rabbit IgG-HRP-conjugated antibody (172-1019, Bio-Rad) at 1:10000. After three more washing steps in PBST, detection was performed via enhanced chemiluminescence (ECL; Amersham, Cytiva).

### Analysis in mouse primary neuron cultures

#### Preparation of murine hippocampal cultures

Animal experiments were approved by the local Animal Care and Use Committee (Regierungspraesidium Tuebingen, Tuebingen, Germany). *Kcnq2^tm1Dgen^/Kcnq2^+^*(*Kcnq2*^-/+^) animals were originally generated by Deltagen and imported from Jackson laboratories (B6.129P2-Kcnq2^tm1Dgen^/J; JAX stock #005830). For murine hippocampal cultures we used E17 embryos from the breeding of 2 heterozygotes. Pregnant females were sacrificed using CO_2_ and embryos were decapitated quickly. Dissected hippocampi were washed three times with 4°C Mg^2+^- and Ca^2+^ free HBSS (PAA Laboratiories, Germany) followed by incubation for 15 min with 2,5 % trypsin. Afterwards, hippocampi were rinsed in DMEM plus 10% fetal bovine serum (Biochrom, Germany), L-Glutamine (Thermo Fisher, USA) and penicillin/streptomycin (Thermo Fisher, USA) to block the trypsin. To obtain single cells the hippocampi were mechanically dissociated by pipetting and passed through a cell strainer (Becton Dickinson, Germany). 150 000 cells in 110 µl were plated on a MEA array (Multi Channel Systems, Germany). After settling down for 4 hours, 1 ml of Neurobasal culture medium (Thermo Fisher,USA) supplemented with B27 (Thermo Fisher, USA), glutamine and penicillin/streptomycin was added (Liu et al. 2019). MEA dishes were coated with poly-D-lysine (5mg in 100 ml of HBSS) at room temperature for 4 hours. After washing with PBS (Thermo Fisher, USA), 10 µl of a laminin coating solution in a concentration of 100 µg/ml was applied onto the electrodes overnight. MEA dishes used had a square grid of 60 planar Ti/TiN electrodes of 30 µm diameter with 200 µm spacing (Multi Channel Systems, Reutlingen, Germany).

#### MEA recordings and analysis

Recordings were performed on days 10 and 21 after plating. Culturing medium was changed one day before the recording (Neurobasal with B27 and glutamine). The recordings of spontaneous activity lasted 2 minutes, after which the dishes were returned to the incubator. The recordings were sampled at 25kHz, visualized and stored using the standard software MC_Rack provided by Multi Channel Systems. For Spike detection we used SPANNER software (RESULT Medical, Germany). Synchronous network activity was analysed by population burst detection using custom-built Matlab software.

### Quantification and statistical analysis

#### Electrophysiological data analysis

The inclusion of cells in the analysis was based on passing the following criteria: RMP<-50mV and firing more than one spike in evoked firing protocols. These criteria were not applied to very immature neurons at DIV 3-4 (NGN2 protocol). A spike was defined as an at least 25 mV positive deviation from the interspike plateau/baseline. The recorded data were analysed by Clampfit (Molecular Devices, USA), Excel (Microsoft, USA) and GraphPad Prism 9 (Dotmatics, USA).

#### GDE data processing

Current signals from up to five minutes long voltage clamp recordings were loaded into MATLAB and processed at their original sampling rate of 100,000 Hz. For low-frequency events, data were band-pass filtered between 0.5 and 5 Hz using a 10,000-order FIR filter, linear trends in the entire recordings were removed and data were smoothed using a moving average of one second. Subsequently, to identify giant depolarizing events (GDEs), we performed peak detection on the one-second smoothed, detrended data. We chose a minimum peak distance of 5 seconds, and an adaptive minimum peak prominence above the neighboring signal amplitude of 0.4 × maximum of signal modulus, but at most 100 pA. To compensate for signal trends on shorter time scales of the order of the interval between peaks, we individually removed any linear trend in each inter-peak interval around each peak.

In order to determine the temporal extent, i.e. start and end, of each GDE, we used an adaptive procedure. We first used a threshold of 0.9 × peak prominence above each peak minimum, if it intersected the signal close to the peak, and otherwise lowered the threshold. We used the two intersections of signal and threshold closest to each peak as candidates for GDE start and end. We additionally determined a second choice of GDE start and end candidates as the sharp kink in the leading edge of the GDE closest to the peak, and the corresponding intersection with se same level after the peak. We finally selected as GDE start and end, whichever of the two candidates was closer to the peak, thus implementing a conservative choice of GDE extent. Individual GDEs were also confirmed by visual inspection. Eventually, GDE profiles were determined by dividing each GDE into ten deciles and calculating signal amplitude averages in each decile, thus accounting for different GDE duration.

In order to assess firing intensity during giant depolarizing events, raw data were additionally band-pass filtered in a high-frequency band between 10 and 100 Hz using a 10,000-order FIR filter. Subsequently, peaks were detected in this high-frequency band that exceeded 10 standard deviations of the signal. Finally, within each giant depolarization event, its spike firing rate was calculated as number of peaks within the GDE divided by its duration.

Current clamp data (Carbamazepine effects on GDE) were divided into 2 frequency windows: 0.05 - 5 Hz for low-frequency events and 10-100 Hz for high-frequency events. The bursts were found using Matlab’s ‘findPeaks’ function with a minimum peak distance of 1 s between 2 peaks, the start and the end were pointed by a signal rise above 2.5 mV. The high-frequency window was analysed by a previously mentioned threshold of 8 standard deviations (Rosa et al. 2020).

#### Statistics

All column analyses data were analysed for normal distribution (D’Agostino-Pearson). Data sets comparing 2 groups were analysed by Welch’s t-test (normal distribution) or Mann-Whitney test (non-normal distribution), in a case of more than 2 studied groups we used one-way ANOVA (normal distribution) or Kruskal-Wallis test (non-normal distribution). For a post-hoc analysis default tests (Holm-Sidak’s and Dunn’s tests) of GraphPad Prism (Dogmatics, USA) were used. For before/after experiments we used paired alternatives of the above-mentioned tests. The evoked firing curves were analysed using two-way ANOVA for repeated measurements of matched values for all respective current injections. The analysis was performed using GraphPad Prism 9 Software (GraphPad). Since keratinocyte-derived lines showed a biphasic action potential firing profile we used a repeated t-test with an acceptable false discovery rate 1% (two-stage step-up method of Benjamini, Krieger and Yekutieli).

### Study approval

Extraction of skin biopsies for the establishment of fibroblasts cultures or hair plucking for the generation of keratinocyte cultures and their use in the generation of induced pluripotent stem cells (iPSCs) were approved by the Ethics Committee of Medical Faculty and the University Hospital Tübingen. All donors signed the consent form.

## Data and software availability

Data and analysis routines are available upon reasonable request to the corresponding authors.

## Author contributions

Conceptualization: F.R., S.M., T.V.W.; Methodology: F.R., He.L., S.K., S.L.; Software: S.T., D.M.; Investigation: F.R., S.K.; Writing-original draft preparation: F.R., S.M., T.V.W., S.T.; Writing-Review and Editing: all authors; Resources: H.L., S.P.; Funding acquisition: H.L., S.M., T.V.W., F.R.; Supervision: T.V.W., S.M., H.L.

## Funding / Acknowledgements

This work was supported by the German Federal Ministry for Education and Research (BMBF): Rare disease networks IonNeurONet (01GM1105A to SM and HL) and Treat-ION (01GM2210A to HL) and via the European Joint Program on Rare Diseases (EJP-RD), TreatKCNQ (01GM2003B to TVW). FR was supported by the DAAD program (91529082) and Endeavour research fellowship program (6093_2017). We thank the patients and their parents for participating in the study.

## Supplementary Figures

**Supplementary Figure 1.**
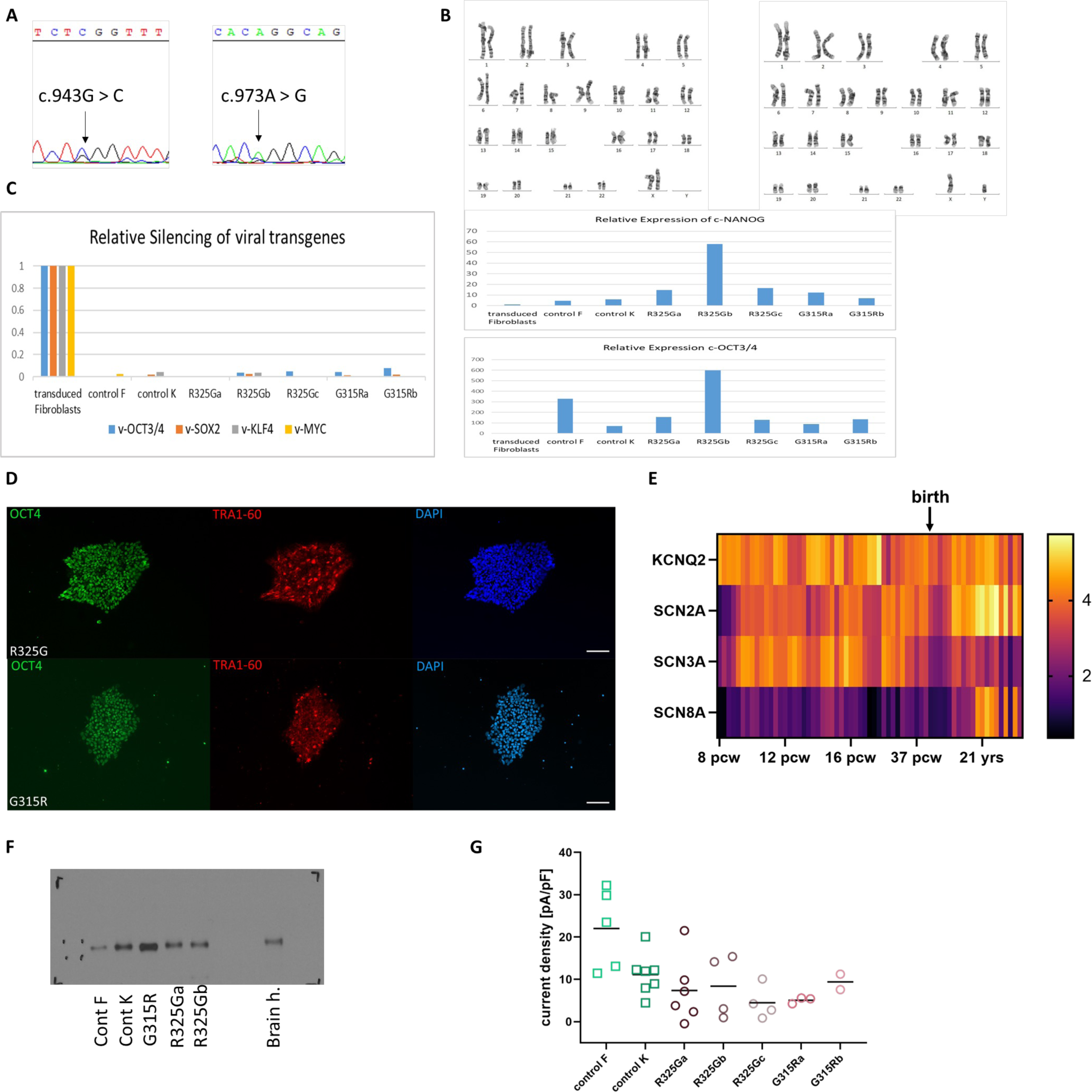
Variants, karyograms, silencing of viral transgenes, expression of pluripotency markers. (A) Chromograms of Sanger sequencing confirming the presence of variants G315R (left) and R325G (right) in corresponding patient lines. DNA was extracted from iPSC-derived neuronal cultures. (B) Karyograms obtained from patient iPSC lines G315R (left) and R325G (right). (C) RT-qPCR experiments showing silencing of viral genes (left), expression of c-NANOG (right top) and expression of c-OCT3/4 (right bottom). (D) Immunostainings in R325G and G315 iPSC lines for OCT4 and TRA1-60. (E) Heat map of the developmental expression of the selected genes according to Allen Brain Atlas. Colour scale of RPKM (Reads Per Kilobase Million) right. Expression in five different brain areas (dorsolateral prefrontal cortex, orbital frontal cortex, primary motor cortex, posterior (caudal) superior temporal cortex, inferolateral temporal cortex) is plotted over development. (F) Western-blot showing the expression of Kv7.2 protein in NGN2 neurons derived from control and patient iPSC lines. Human brain lysate obtained from brain surgery was used as a positive control. (G) Scatter plot of the M-current density at the clamped voltage of 25 mV for all 7 established iPSC-lines (NGN-2 protocol; control F, control K, R325G with indices a,b and c and G315R with indices a and b).

**Supplementary Figure 2.**
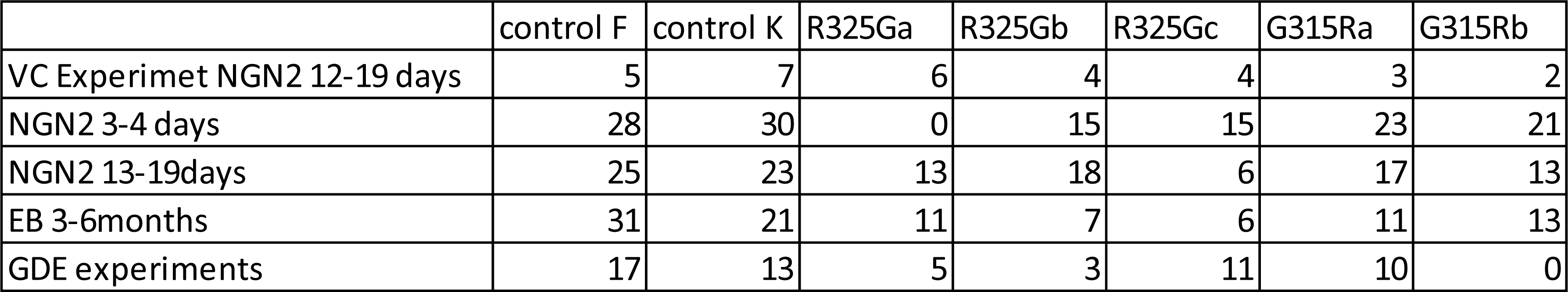
Allocation of iPSC lines to different experiments.

**Supplementary Figure 3.**
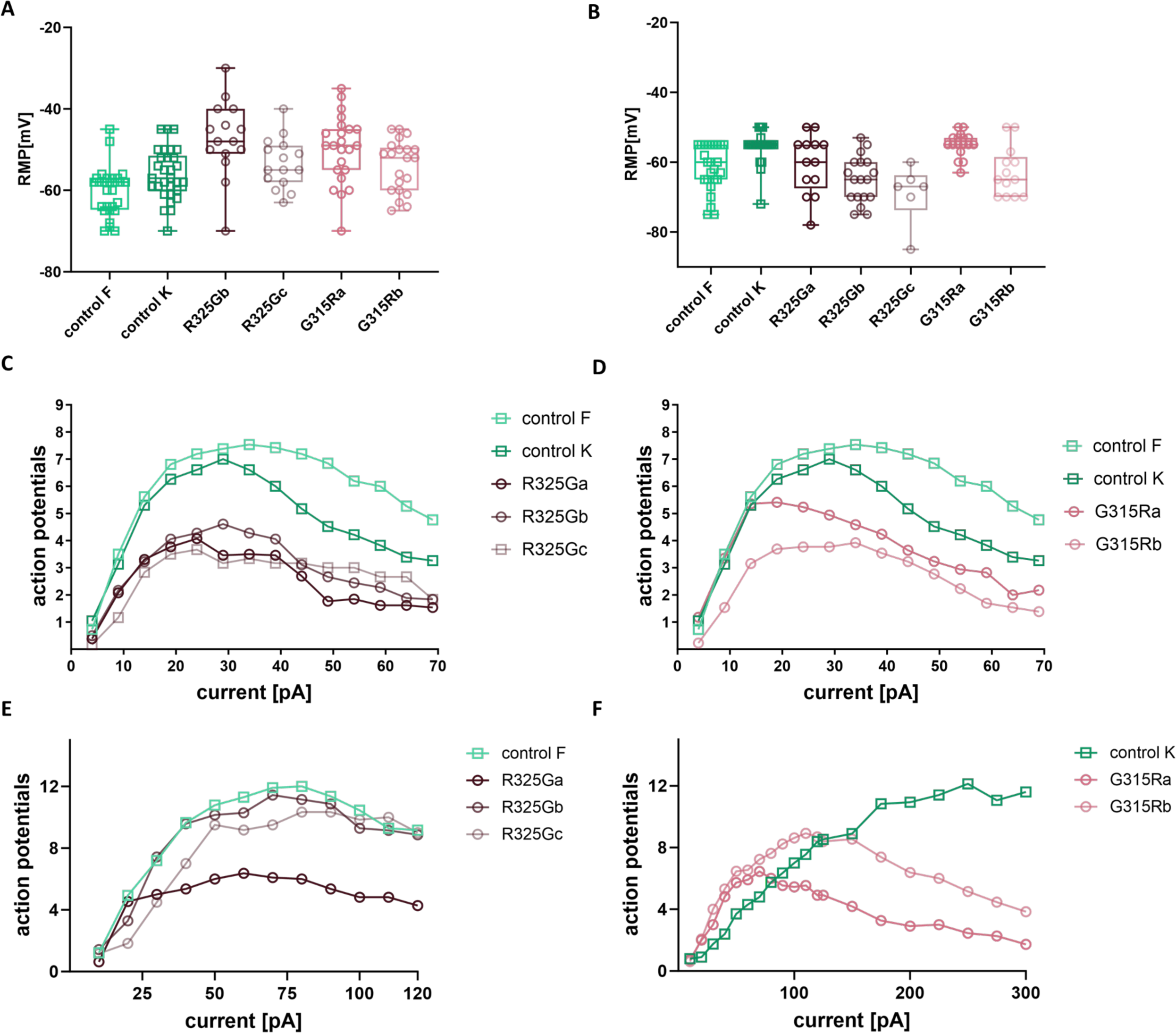
RMP and input-output curves for individual iPSC lines. (A) RMP of NGN2 differentiated iPSC lines recorded between the 3^rd^ and 4^th^ day after start of differentiation. (B) RMP of NGN2 differentiated iPSC lines recorded between the 14^th^ and 19^th^ day after start of differentiation. (C) NGN2 Input-output curve of the evoked firing experiment for controls and all particular R325G lines. (D) NGN2 Input-output curve of the evoked firing experiment for controls and all particular G315R lines. (E) EB Input-output curve of the evoked firing experiment for controls and all particular R325G lines. (F) EB Input-output curve of the evoked firing experiment for controls and all particular G315R lines. Number of recorded cells is stated in Suppl. Fig. 2

**Supplementary Figure 4.**
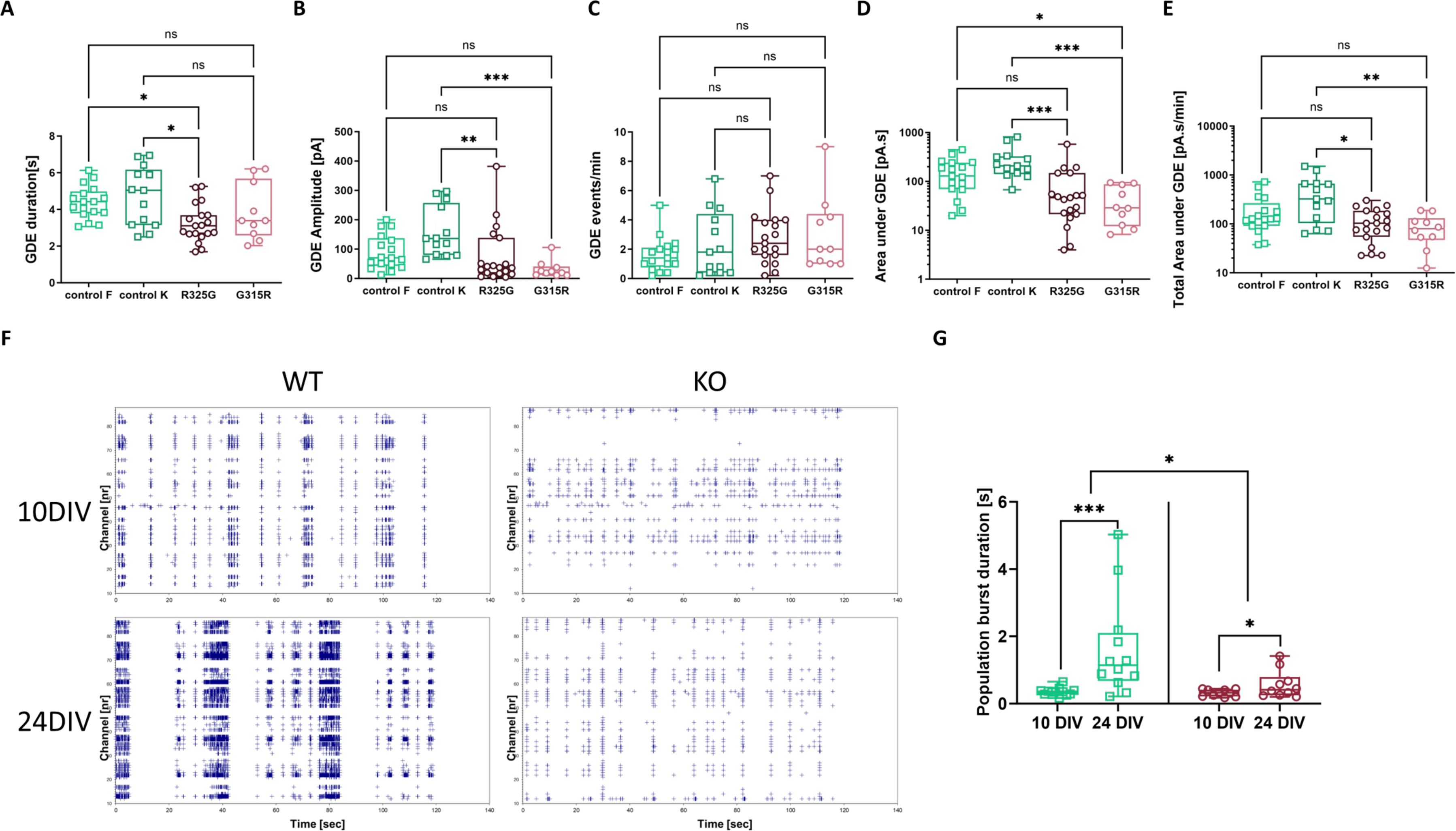
GDE properties according to genotype and comparison to bursting properties of WT and *Kcnq2* knock-out murine primary neuron cultures. (A) GDE duration for non-pooled control and patient lines. Kruskal-Wallis test P = 0.0122; Dunn’s multiple comparison test: control F vs. R325G= 0.0278; control F vs. G315R> 0.9999; control K vs. R325G=0.0377; control K vs. G315R>0.9999; control F 4.38 ± 0.22 s, control K 4.72 ± 0.45 s, R325G 3.24 ± 0.24 s, G315R 3.87 ± 0.49 s; n(control F)=17, n(control K)=13, n(R325G)=19, n(G315R)=10. (B) Peak amplitude of GDEs for non-pooled control and patient lines. Kruskal-Wallis test P = 0.0002 Dunn’s multiple comparison test: control F vs. R325G=0.7728; control F vs. G315R= 0.0505; control K vs. R325G=0.0051; control K vs. G315R=0.0002; control F 88.97 ± 14.24 pA, control K 159.1 ± 23.74 pA, R325G 76.22 ± 22.31 pA, G315R 31.37 ± 9.09 pA; (C) Number of GDE events in one minute for non-pooled control and patient lines. Kruskal-Wallis test P = 0.2130; Dunn’s multiple comparison test: control F vs. R325G=0.3624; control F vs. G315R= 0.7545; control K vs. R325G>0.9999; control K vs. G315R>0.9999; control F 1.61 ± 0.28 GDE/min, control K 2.35 ± 0.61 GDE/min, R325G 2.77 ± 0.41 GDE/min, G315R 3.02 ± 0.81 GDE/min; (D) Area under an average GDE event for non-pooled control and patient lines. Y-axis logarithmic. Kruskal-Wallis test P < 0.0001; Dunn’s multiple comparison test: control F vs. R325G= 0.0844; control F vs. G315R=0.0280; control K vs. R325G=0.0009; control K vs. G315R=0.0004; control F 162.7 ± 28.97 pA.s, control K 285.9 ± 61.84 pA.s, R325G 88.88 ± 30.62 pA.s, G315R 43.65 ± 11.09 pA.s (E) Total area under GDE event per one minute for non-pooled control and patient lines. Y-axis logarithmic. Kruskal-Wallis test P = 0.0050; Dunn’s multiple comparison test: control F vs. R325G= 0.9584; control F vs. G315R= 0.3583; control K vs. R325G= 0.0214; control K vs. G315R=0.0094; control F 203.4 ± 46.16 pA.s/min, control K 495.5 ± 126.6 pA.s/min, R325G 117.5 ± 18.69 pA.s,/min G315R 89.96 ± 18.91 pA.s/min (F) Spike raster plots obtained from MEA recordings in WT and *Kcnq2* knock-out cultures at days in vitro (DIV) 10 (top) and DIV 24 (bottom). (G) Comparison of population burst duration between *Kcnq2* WT and KO cultures between DIV10 and DIV24. WT DIV10: 0.36 ± 0.04 s, WT DIV24: 1.62 ± 0.43 s; Ratio paired t-test P=0.0009, t=4.525; n(WT)=12. KO DIV10: 0.319 ± 0.033 s, DIV24: 0.568 ± 0.131 s. Ratio paired t-test P=0.0431, t=2.353; n(KO)=10. Ratios of WT DIV24/DIV10 vs KO DIV24/DIV10 - WT: 4.82 ± 1.33, KO: 1.79 ± 0.37; Mann-Whitney test P=0.0426, U=29; n(WT)=12, n(KO)=10. For multiple comparisons we used Dunn’s test. P<0.05: *; P< 0.01: **, P< 0.001: ***, P<0.0001: ****

**Supplementary Figure 5.**
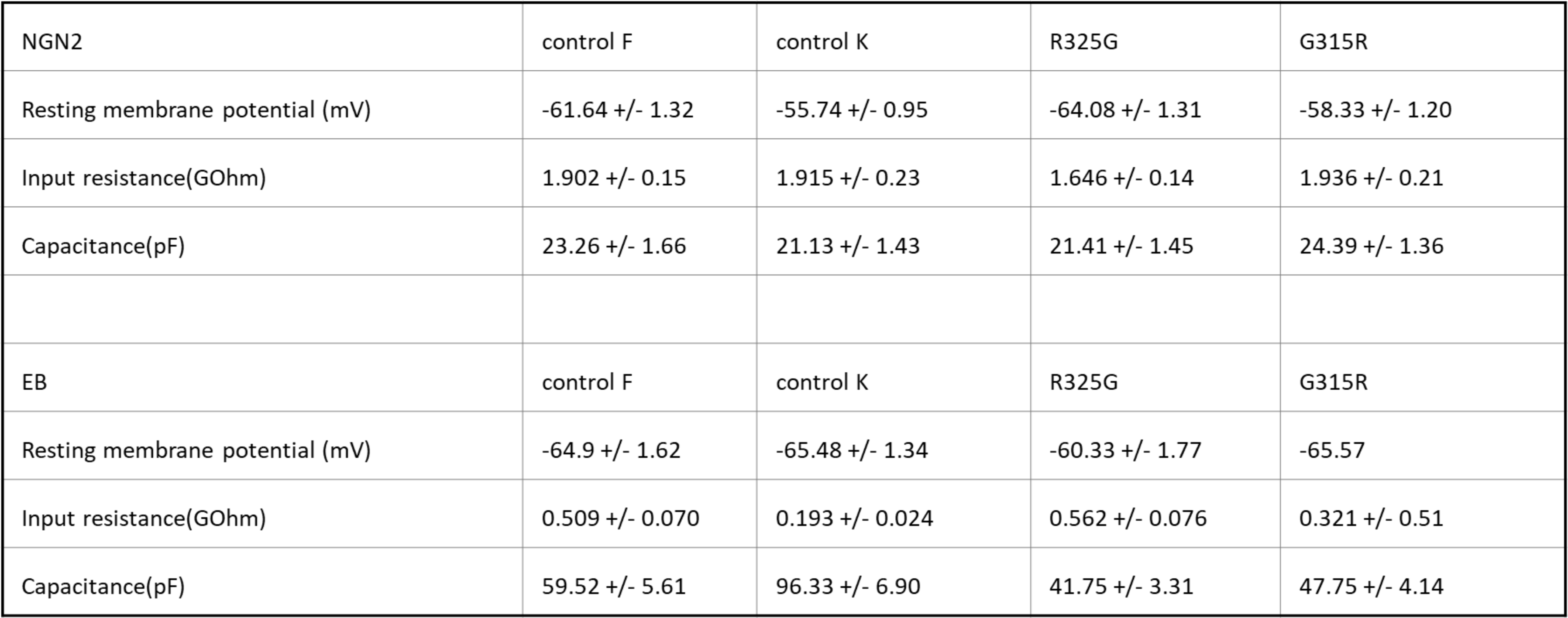
Table of passive membrane properties.

**Supplementary Figure 6.**
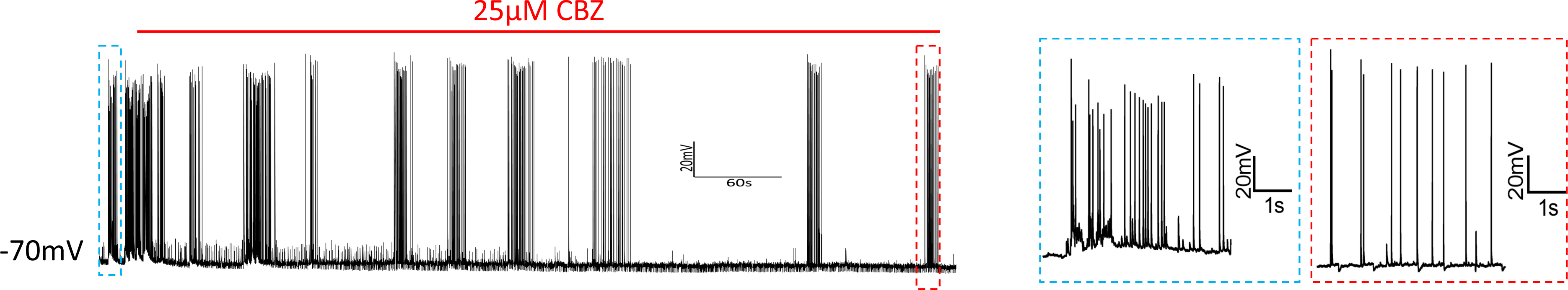
Exemplified conversion of GDE bursting to single action potential spiking by carbamazepine. 10 min spontaneous activity recorded in current clamp mode (left). Blue: a magnified picture of firing before CBZ application; red: a magnified picture of firing after CBZ application.

